# Spreading depolarization, not seizure activity, drives immediate early gene expression in mouse cortex after electroconvulsive stimulation

**DOI:** 10.64898/2026.04.14.718362

**Authors:** Hugo J. Ladret, Leonardo Lupori, Lorenzo Sieni, Eduard Stroukov, Takahiro Kanamori, Sarah Ulrich, Else Schneider, Gunnar Deuring, Annette B. Brühl, Georg B. Keller

## Abstract

Electroconvulsive therapy (ECT) is a highly effective treatment for several psychiatric disorders, though its biological mechanisms remain unclear. ECT’s therapeutic action has traditionally been attributed to the generalized seizure it induces. However, this view is challenged by the recent finding that electroconvulsive stimulation (ECS) can trigger a spreading depolarization (SD) in cortex. Because SD drives massive molecular changes, including expression of the plasticity-related immediate early gene Fos, we hypothesized that SD could be a key mediator of ECT’s plasticity-inducing effects. We recorded the neuronal response to ECS in mouse dorsal cortex and observed similar activity to that of patients undergoing ECT. SD, rather than the seizure activity, accounted for increased Fos expression in mouse dorsal cortex. SD was also associated with EEG features that predict positive treatment outcome of ECT in patients. Our results suggest that the therapeutic efficacy of ECT may be mediated by SD and imply that a currently unmonitored neurophysiological event may serve as a relevant biomarker to predict and optimize ECT outcomes.

## INTRODUCTION

Electroconvulsive therapy (ECT) is one of the most potent and rapidly acting interventions in clinical psychiatry. It demonstrates substantial efficacy in treating severe and often treatment-resistant conditions, including major depressive disorder (Trifu et al., 2021), bipolar disorders (Tor et al., 2021), catatonia (Perugi et al., 2017), mania (Martin et al., 2025) and schizophrenia (Grover et al., 2019). ECT frequently achieves positive therapeutic outcomes in terms of remission rates and speed of response that surpass standard pharmacotherapies, making it indispensable in urgent clinical situations such as high suicide risk or severe catatonia (Baghai and Möller, 2008). ECT is also one of the few treatment options available to a considerable proportion of patients who do not respond to antipsychotic drugs (Haddad and Correll, 2018). Despite its established clinical utility and decades of refinement aimed at enhancing safety and tolerability, the fundamental neurobiological mechanisms driving ECT’s therapeutic effects remain poorly understood.

There is a multitude of biomarkers affected by ECT that could underlie its mode of action as a psychiatric treatment (Singh and Kar, 2017). These include, non-exhaustively, the modulation of various neurotransmitter systems (Baldinger et al., 2014), alterations in regional cerebral blood flow (Takano et al., 2011) and metabolism (McCormick et al., 2007), changes in functional brain network connectivity (Ulrich et al., 2025), and the promotion of neuroplasticity (Bouckaert et al., 2014; Chen et al., 2018; Joshi et al., 2016). Understanding how these effects contribute to therapeutic outcomes is a significant challenge. We suspect that a critical gap in our knowledge lies in the early cascade of events triggered by an ECT treatment.

ECT, as it is used today, is targeted at the induction of a controlled seizure under general anesthesia and muscle relaxation through an electroconvulsive stimulation (ECS). ECS-triggered seizures specifically refer to highly synchronized brain-wide oscillatory activity, which typically follows a three-phase progression (Brumback and Staton, 1982) consisting of an immediate 14-22 Hz activity (phase I), which evolves into arrhythmic poly-spike patterns (phase II) and is then rapidly, within seconds, followed by a sustained low-frequency oscillatory activity at 2-4 Hz (phase III). Phase III activity often occurs without any ictal spikes that are characteristic of epileptic seizures (Brumback and Staton, 1982) and is of lower amplitude following ECT compared to the late phase of seizures observed in epilepsy (Pottkämper et al., 2021). The phase III activity is referred to as seizure in the ECT literature. Illustrating the difference in meaning of the term seizure in the ECT literature and the epilepsy literature is also the fact that the post-seizure neuronal silencing of activity is a marker of treatment success in ECS, while it is associated with sudden unexpected death in epilepsy (Krystal et al., 1995; Sowers et al., 2013). To try to disambiguate this, we will refer to the ECS-induced seizure as phase III activity in the description of our results, but consistent with previous usage as seizure when talking about prior ECT work.

Historically, the generalized ECS-induced seizure was considered to be the essential therapeutic component of the treatment (Accornero, 1988; Ulett et al., 1956). However, the occurrence and properties of seizures are only partially predictive of therapeutic outcomes (Perera et al., 2004; Sackeim et al., 1991). Similarly, the clinical efficacy of alternative stimulation methods, such as low amplitude ECT or magnetic seizure therapy, demonstrates that robust outcomes can be achieved with lower field strengths (Peterchev et al., 2015). This discrepancy suggests that electric field strength is unlikely to be the primary determinant of therapeutic success (Deng et al., 2024), indicating that ECT’s efficacy hinges on biological mechanisms beyond the mere presence of a seizure or the magnitude of the electrical stimulus.

A secondary neurophysiological event frequently triggered by seizures is spreading depolarization (SD). It has been speculated that SD is an endogenous mechanism of the brain used for seizure termination (Dreier et al., 2012; Kelly et al., 1999). An SD can also be triggered by direct electrical stimulation of the cortex (Fregni et al., 2007; Leao, 1944), and, consistent with this, recent evidence revealed the existence of SD-like calcium events following ECS with on-skull electrodes in mice and likely following ECT in patients (Rosenthal et al., 2025). This discovery necessitates a critical re-evaluation of the traditional seizure-centric view of the effects of ECT. The profound neurophysiological disruption inherent to SD may potentially act as a trigger for several of the neurobiological alterations driven by ECT.

SD is characterized as a slowly propagating wave (typically 2–5 mm/min) of near-complete and sustained depolarization affecting both neurons and glial cells (Seidel et al., 2016). This wave of intense depolarization is invariably followed by a period of marked suppression of spontaneous and evoked electrophysiological activity that lasts several minutes and can persist for up to an hour (Sawant-Pokam et al., 2017). The wavefront of the SD is a massive transmembrane ionic flux (Somjen, 2001) that is accompanied by cellular swelling due to water movement, blood flow changes resulting in local hypoxia, and structural changes to dendritic processes (Takano et al., 2007). An SD can be triggered by a large efflux of potassium ions into the extracellular space (Brinley et al., 1960) that causes a positive feedback loop via glutamate release, which propagates the wavefront through the activation of NMDA receptors (Zhou et al., 2013). Concurrently, there is an influx of sodium, chloride and calcium ions into cells (Pietrobon and Moskowitz, 2014). Disturbed calcium signaling is associated with a host of neuropsychiatric disorders treated by ECT (Nanou and Catterall, 2018), which makes SD-driven calcium influxes a likely key contributor to neuronal plasticity effects driven by ECT (Dehbandi et al., 2008; Wernsmann et al., 2006).

To assess whether SD drives plasticity in the brain, we measured the expression of the protein Fos following ECS. Fos is the product of an immediate early gene transcribed following high levels of intracellular calcium influx (Chung, 2015). Fos is often used as a marker of recent neuronal activity, and its expression can be driven by strong artificial activation of neurons (Tyssowski et al., 2018). However, in cortex, Fos expression correlates only weakly with neuronal activity (Mahringer et al., 2022). Conversely, Fos is required for several forms of synaptic plasticity and long-term memory (Fleischmann et al., 2003; Watanabe et al., 1996), and in hippocampus marks neurons that exhibit learning-related activity changes (Mahringer et al., 2019). Consistent with this, early experiments using natural sensory stimulation found that the first exposure to light after dark rearing, a stimulus that triggers experience-dependent plasticity in visual cortex, strongly induced Fos, and that this induction was larger during the critical period than in adulthood (Mower, 1994; Rosen et al., 1992). Thus, Fos expression is likely better interpreted as a readout of a transcriptional program of plasticity than as a marker of recent activity. Both ECS (Calais et al., 2013; Winston et al., 1990) and SD (Ingvardsen et al., 1997; Iqbal Chowdhury et al., 2003) are known to induce Fos expression in the brain. Given that the seizure is used as a clinical marker for treatment efficacy, the key question is, what fraction of the ECS-induced plasticity is caused by the seizure and how much by the SD.

To address this, we sought to dissociate the contributions of SD and phase III activity to ECS-induced Fos expression, hypothesizing that the calcium dynamics specific to SD underlie the plasticity-relevant effects of ECT. Consistent with previous work (Rosenthal et al., 2025), we observed a slow traveling calcium event with the characteristics of an SD. This event accounted for the ECS-driven Fos expression in the dorsal cortex. Furthermore, we show that EEG metrics known to predict positive therapeutic outcomes of ECT are correlated with the occurrence of SD in mice. Our findings support the hypothesis that SD, rather than other components of the neuronal response to ECS, is the primary driver of ECS-driven plasticity and, by extension, might drive the therapeutic benefits of ECT.

## RESULTS

### ECS triggers similar neuronal activity in both mice and humans

We first asked how well ECS in mice recapitulates EEG signatures of ECS in humans. To do this, we measured neuronal responses following ECS in mouse dorsal cortex, using concurrent widefield calcium imaging and EEG recordings (**Methods**). We expressed a genetically encoded calcium indicator brain-wide by retro-orbital deposit of an AAV-PHP.eB-Ef1α-GCaMP6s-WPRE and performed concurrent EEG recordings using a semitransparent 30-channel EEG electrode (**Figure 1A**). During ECS, mice were anesthetized with isoflurane (2%) and myorelaxed with succinylcholine chloride (1 mg/kg). ECS was delivered through biauricular electrodes using a commercial ECS unit. Stimulation parameters were set within a range that reliably evoked tonic-clonic seizures in mice (**Methods**).

**Figure 1.**
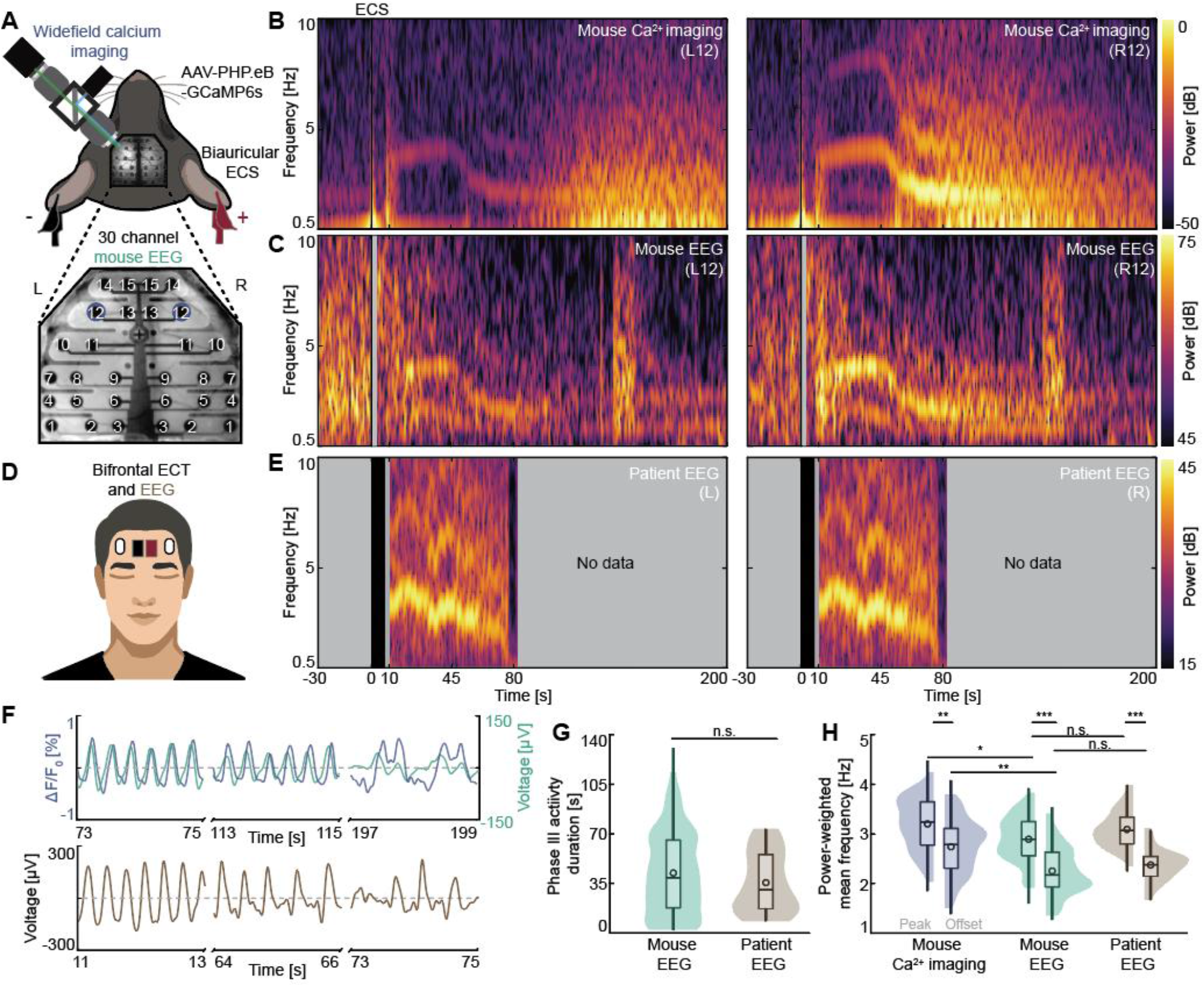
Electroconvulsive stimulation (ECS) triggers similar slow neuronal activity in mice and humans. (**A**) Top: Schematic of mouse ECS setup. We performed concurrent widefield calcium imaging and EEG in the mouse dorsal cortex during biauricular ECS. Bottom: Layout of the EEG probe. (**B**) Spectrograms of two widefield calcium imaging ROIs surrounding EEG electrodes L12 and R12 (as highlighted in **A**). Duration of the ECS is marked by a black bar. (**C**) Spectrograms of the EEG recorded concurrently with the widefield calcium data in **B**. Time during which EEG is not recorded due to a stimulation artifact is marked by gray shading. (**D**) Schematic of patient ECT setup. EEG recordings were performed using two frontal electrodes after bifrontal ECT in patients. (**E**) Spectrograms of EEG recorded following ECT in a patient. Patient EEG recordings are started following ECT and terminated when the phase III activity stops. Gray shading marks times without EEG recording. (**F**) Top: Phase III activity observed in early, mid, and late phases after ECS in an example mouse widefield calcium ROI (blue) and in the concurrently recorded EEG (green). Bottom: Phase III activity observed in an example patient EEG recording. (**G**) Phase III activity duration for mouse and patient EEG recordings. The central box represents the interquartile range (IQR), spanning from the first (Q1) to the third (Q3) quartiles. Whiskers extend to data points not considered outliers (**Methods**). A central horizontal line indicates the median and an open circle the mean of the data distribution. (**H**) Power-weighted mean frequency at peak and offset of phase III activity for mouse widefield, mouse EEG and patient EEG recordings. Here and elsewhere: n.s.: not significant, *: p < 0.05, **: p < 0.01, ***: p < 0.001. See **Table S1** for all information on statistical testing.

Across the parameter range used, ECS triggered an immediate and brief calcium response across the entire dorsal cortex (**Figure S1A**). The response onset occurred with a latency of less than 10 ms (within the first frame after stimulation at 100 Hz acquisition), and calcium activity continued to increase until the end of the ECS (**Figure S1B**). In the EEG data (**Figure S1C**) we were blind to this direct response, as we disconnected the EEG from the data acquisition system during ECS to prevent electrical arcing between electrodes (**Methods**). We speculate that the direct ECS response observed in calcium imaging is driven by spiking activity in the cortical neurons, given that the electrical field generated by the ECS is several times higher than the neuronal activation threshold (Deng et al., 2011). Following the direct calcium response to ECS, mean calcium activity decreased and returned to baseline only slowly (**Figure S1B**).

Following ECT, the combined median duration of phases I and II activity is 9 s in patients (Hogan et al., 2019). Consistent with this and previous reports (Murakami et al., 2008), we observed a phase III slow oscillatory activity in the delta to low theta range (1 to 5 Hz) in mice that emerged about 10 s after the ECS, both in calcium activity and EEG recordings (**Figure 1B-C, S2**). This phase III activity was strikingly similar to that observed in patients undergoing ECT (**Figure 1D-E**) (Stuiver et al., 2026), and was present in the EEG recording of all patients analyzed here. Given that ECT stimulation parameters are chosen to increase the likelihood of phase III activity, which in turn is used as a biomarker for treatment outcome (Sackeim et al., 1991), this is perhaps not surprising.

The phase III activity followed a similar pattern in both mice and patients and typically exhibited a strong sinusoidal shape in the early stage that desynchronized over time (**Figure 1F**). In mice, phase III activity lasted 42.3 ± 22.1 s (mean ± std), and we found no evidence of a difference in activity duration between mice and patients (**Figure 1G**). These values are consistent with previous estimates of the duration of phase III activity in patients (Hogan et al., 2019). Interestingly, the frequencies of the activity were also very similar in mice and patient EEG, and both tended to decrease with time (**Figure 1H**). The estimate of the activity frequency in the mouse calcium imaging data was systematically higher relative to the concurrent EEG data (**Figure 1H**) but exhibited the same decreasing trend. We suspect this difference is driven by differential contributions to the two signals from non-phase III related neuronal activity and noise sources.

### ECS can trigger a traveling calcium event

We found that ECS could also trigger a traveling calcium event, as previously described (Rosenthal et al., 2025). Based on several of its characteristics, it has been proposed that this calcium event is an SD (Rosenthal et al., 2025). While we agree with this interpretation, we will refer to it as a calcium event in the results section here to avoid forgoing the conclusion. To monitor the propagation of these events, we recorded the calcium activity across the entire dorsal cortex for 3 to 5 minutes following ECS (**Figure 2A**). They manifested as a significant increase in calcium fluorescence that propagated across the dorsal cortex (**Figure 2B**) at a speed of 0.083 ± 0.009 mm/s (mean ± std) (**Figure 2C**), consistent with previously reported SD propagation speeds in mice (Enger et al., 2015). Calcium events exhibited maximum fluorescence intensities that exceeded even those observed during the direct ECS response (**Figure 2D**). Once initiated, these events spread across an entire dorsal cortex hemisphere over the course of several minutes (**Figure 2E**). In line with the refractory dynamics of SD, the event exhibited a non-repeating trajectory, traversing each part of dorsal cortex only once. The lateralization of the events exhibited a certain stochasticity and calcium events were not always symmetrically present in both hemispheres. We frequently observed cases in which the events were unilateral and constrained to only left or right cortical hemispheres (**Figure 2F-G**).

**Figure 2.**
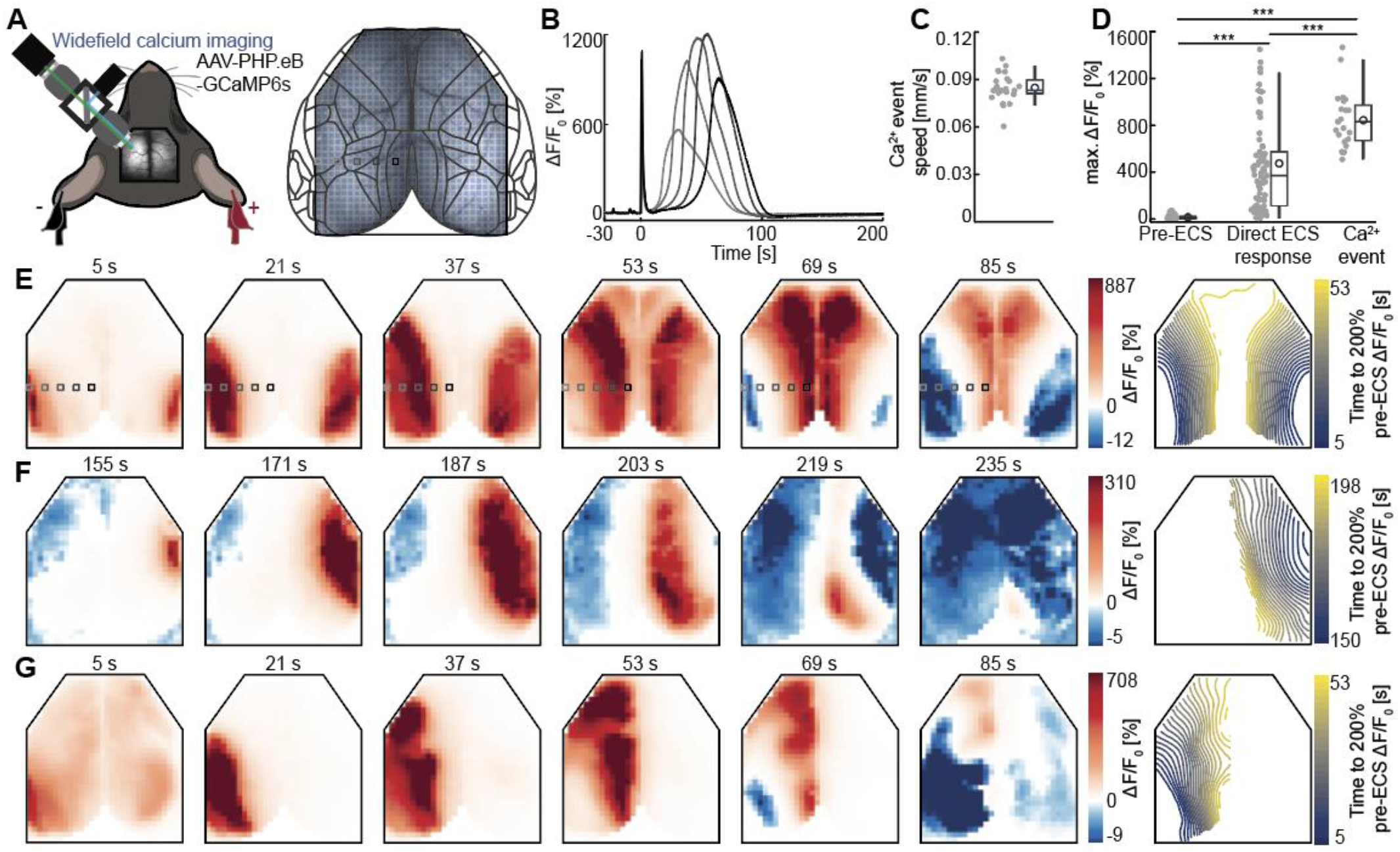
ECS can trigger a traveling calcium event. (**A**) Left: Schematic of mouse ECS setup without concurrent EEG recordings. Right: Widefield recordings of mouse dorsal cortex were parcellated into 818 ROIs, each of which was assigned to a cortical region according to the Allen Brain Atlas taxonomy. The ROIs highlighted correspond to the calcium activity shown in **B** and **E**. (**B**) Example of calcium activity in ROIs highlighted in **A** during the traversal of a calcium event. (**C**) Propagation speed of calcium events (**Methods**). Each data point is the median speed across one calcium event. (**D**) Maximum calcium activity throughout the dorsal cortex during pre-ECS, direct ECS response and calcium event periods. Two direct ECS responses are outside the plot range and not shown (1707 % and 2374 % ΔF/F0). (**E**) Left: Low-passed (< 0.5 Hz) calcium activity during an example ECS session triggering a calcium event over both hemispheres. Each panel shows the average calcium activity over a 10 ms window starting at the time indicated above the panel. Right: Isochrones (spaced at two-second increments) indicating the time at which each region reaches 200% of its baseline fluorescence. (**F**) As in **E**, but for a unilateral calcium event over the right hemisphere following a 150 s delay post-ECS. (**G**) As in **E**, but for a unilateral calcium event over the left hemisphere.

In about one third of cases, calcium events appeared bilaterally, while in the remaining two thirds the events were confined to one hemisphere only (**Figure 3A, Video S1**). The majority of unilateral events occurred in the right hemisphere, and across all events, most first appeared on the right edge of the cranial window (**Figure 3B**). This right hemisphere bias was likely driven by the ECS polarity, as we always placed the positive pole of stimulation on the right ear (**Figure 1A**). Calcium events were the only type of response to ECS that occurred unilaterally (**Figure 3C**). While all calcium events co-occurred with phase III activity, just over half of the ECS sessions triggering phase III activity also resulted in a calcium event (**Figure 3D**). The likelihood of triggering a calcium event has been shown to be a function of the stimulation current and frequency used of the ECS (Rosenthal et al., 2025). To investigate how the stimulation parameters control the appearance of phase III activity and calcium events, we tested stimulation frequencies between 25 and 100 Hz and stimulation currents between 15 and 75 mA for a total charge delivery of between 0.187 and 3.750 mC. With most combinations of parameters, we observed a non-zero likelihood of triggering phase III activity (**Figure 3E**). The threshold for triggering calcium events was systematically higher and appeared to be a function of both stimulation frequency and current (**Figure 3E**). Based on this, we hypothesized that total charge delivery might be a good predictor of the occurrence of calcium events. A linear regression of total stimulation charge against the likelihood of triggering a calcium event showed a significant correlation (**Figure 3F**). Stimulation frequency and stimulation current also exhibited a positive but nonsignificant correlation with the likelihood of triggering a calcium event (**Figure S3A**). A multivariate linear regression confirmed that the interaction term of current and frequency was significant and non-zero (0.015 % / (mA Hz)). We found a similar relationship between the total stimulation charge and the probability of triggering phase III activity (**Figure 3G, S3B**). Consistent with the hypothesis that the calcium event exhibits all-or-none dynamics, we found no evidence of changes in propagation speed, total duration, or delay to onset of calcium events as a function of total stimulation charge (**Figure S3C-E**). We also observed that the total ECS charge correlated with the intensity of the direct ECS response (**Figure S4A**). This direct response, in turn, predicted the hemisphere of the calcium event in 75% of the unilateral cases (**Figure S4B**), supporting our hypothesis that the ECS stimulation parameters can control the occurrence of calcium events.

**Figure 3.**
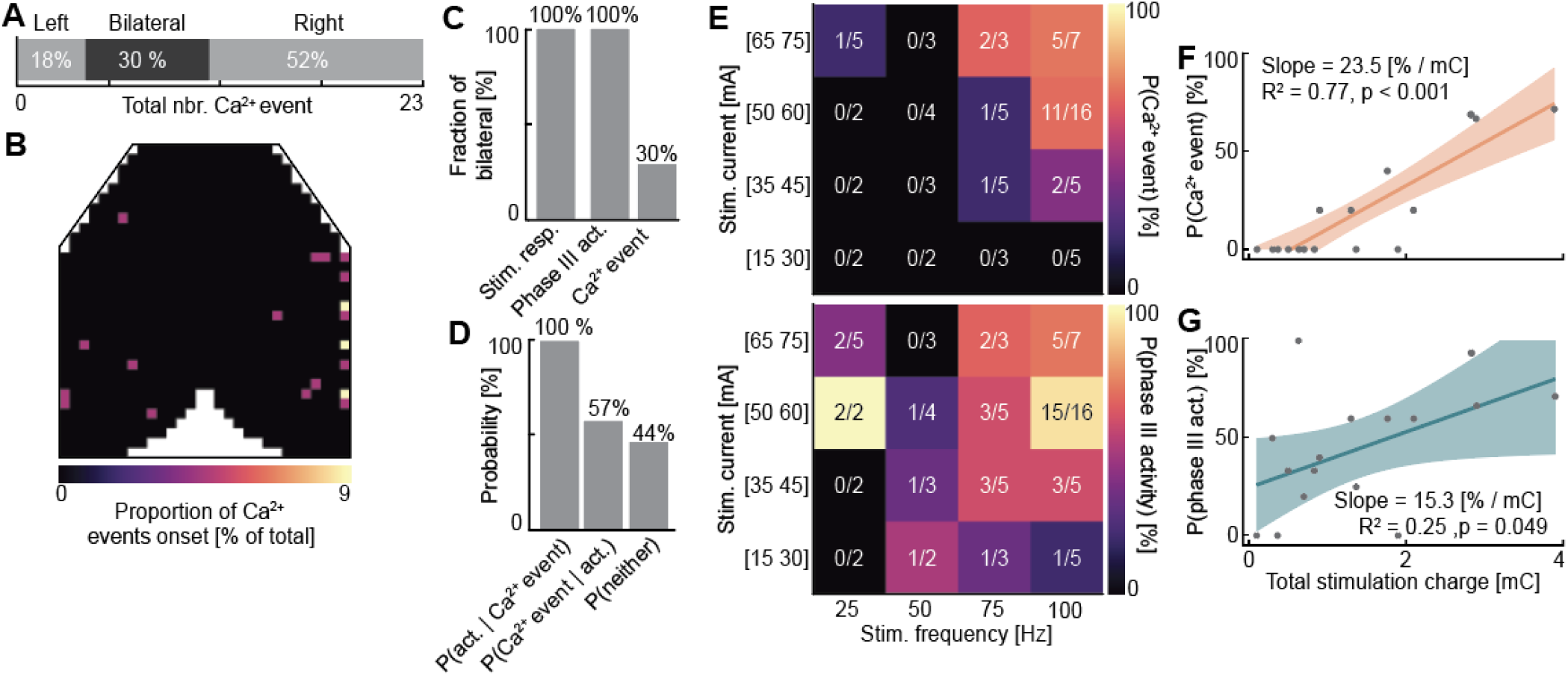
Calcium events are triggered preferentially by higher total ECS charge. (**A**) Probability distribution of the laterality of calcium events. (**B**) Spatial distribution of calcium event detection sites. For bilateral calcium events, only the hemisphere where the calcium event was detected earliest is considered as the detection site. (**C**) Fraction of stimulation responses, phase III activity and calcium events that were bilateral. (**D**) Probability of phase III activity being observed if a calcium event is triggered (P(act. | calcium event)), of a calcium event being observed if a phase III activity is triggered (P(calcium event | act.)), and of neither being observed (P(neither)). (**E**) ECS parameter map of the likelihood of triggering a calcium event (top) or a phase III activity (bottom). (**F**) Probability of triggering a calcium event as a function of total ECS stimulation charge. The solid line represents the linear fit of the data; the shaded region represents 95% confidence interval on the fit. (**G**) As in F, but for the probability of triggering a phase III activity.

### The calcium event exhibits hallmark features of SD

The mode of propagation, speed and fluorescence intensity support the hypothesis that the traveling calcium event is an SD. This is consistent with previous work (Rosenthal et al., 2025), and makes a number of directly testable predictions. First, an SD changes the local ionic homeostasis and is accompanied by hemodynamic constriction (Takano et al., 2007). To test for hemodynamic effects triggered by ECS, we performed widefield imaging experiments in a cohort of mice in which we expressed enhanced green fluorescent protein using an AAV (EGFP; AAV-PHP.eB-Ef1α-EGFP-WPRE). In widefield imaging, the EGFP fluorescence signal depends on pH and overlying blood volume. The latter is a phenomenon called hemodynamic occlusion, whereby decreased blood vessel diameter increases the EGFP signal (Ma et al., 2016; Yogesh et al., 2025). During the ECS, we observed an immediate increase in the EGFP signal that lasted for a couple of seconds and is consistent with acute vasoconstriction (**Figure S5A**). We also observed traveling signals in the EGFP recordings with spatial and temporal characteristics that resembled the calcium event (**Figure S5B**). The amplitude of these events was smaller than that observed in calcium imaging (**Figure S5C**), but they propagated at a speed that did not differ significantly from that observed in calcium events (0.096 ± 0.02 mm/s, **Figure S5D**). This would be consistent with either vasoconstriction or an increase in intracellular pH. SD is associated with vasoconstriction (Takano et al., 2007), but typically only a brief alkalization over the course of seconds followed by a more prolonged acidification of the intracellular domain over the course of minutes (Sun et al., 2011). However, given that an estimated alkalization of less than 0.5 in pH would translate to less than a 10% ΔF/F_0_ change in EGFP fluorescence (Kneen et al., 1998), we suspect most of the change we observe is driven by hemodynamics.

A second feature of SD is the propagation through extracellular diffusion (Enger et al., 2015), rather than through electrical activity. Increases in intracellular calcium levels in neurons are typically associated with increases in spiking activity. An SD, however, is associated only with brief increases in spiking activity at the onset (Herreras et al., 1994; Nasretdinov et al., 2023). To visualize the propagation of the calcium event in the tissue, we used two-photon microscopy to observe the wavefront at cellular resolution. We expressed a genetically encoded calcium indicator (AAV2/9-Ef1α-GCaMP6f-WPRE) in visual cortex and measured responses to ECS in layer 2/3 neurons (**Figure 4A, Video S2**). Consistent with a diffusion-limited process, we found that the calcium event was a sharp wavefront of significant increase in fluorescence, propagating across the entire field of view (**Figure 4B**) at a speed of 0.079 ± 0.0174 mm/s (mean ± std). This aligns with our speed estimates based on widefield calcium imaging. Combined with the lack of propagation along inter-hemispheric or other known anatomical projections (**Figure 2E-G**), this supports the idea that the calcium events are propagating through a diffusion-limited process, rather than synaptic activity, consistent with an SD.

**Figure 4.**
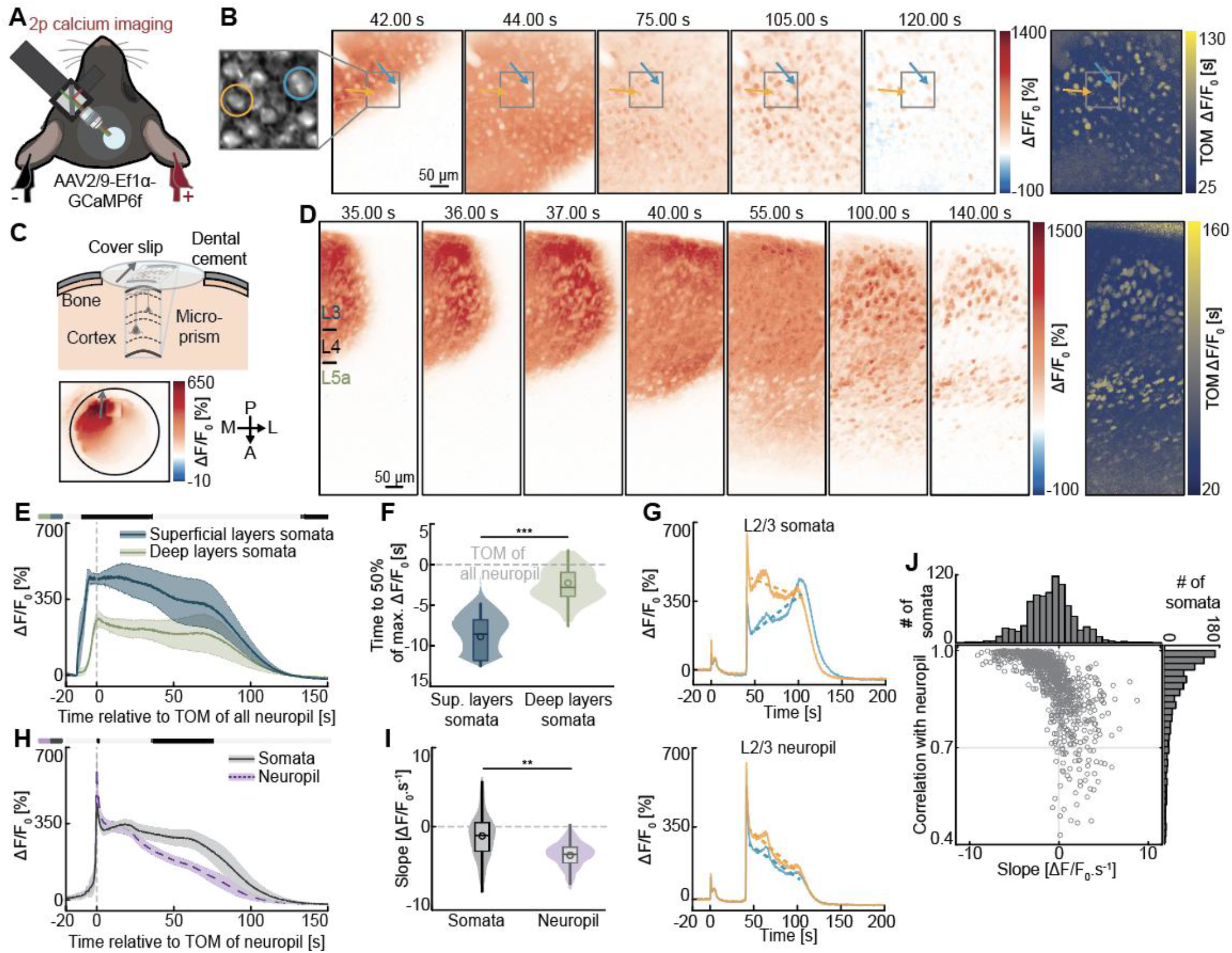
Calcium events first spread in supragranular layers and elicit long-lasting calcium increase in cells. (**A**) Schematic of the experimental setup. We performed two-photon (2p) calcium imaging in the right primary visual cortex during ECS. (**B**) 2p calcium activity during an example ECS session triggering a calcium event. Time of maximum (TOM) is shown on the right. Each panel shows the average calcium fluorescence over a 500 ms window starting at the time indicated above the panel. Gray inset highlights two example somata whose calcium signals are shown in **G**. (**C**) Top: Schematic of the micro-prism implanted below the imaging window to allow for simultaneous recording in all cortical layers (see **Methods**). Bottom: Widefield imaging of the implant 18 seconds after ECS, confirming the prism does not obstruct the calcium event propagation. A gray arrow indicates the orientation of the prism. (**D**) As in **B**, but imaged through a prism implant. Cortical layers 3, 4, and 5a (L3, L4, L5a) are defined based on soma densities (**Methods**). (**E**) Calcium fluorescence of superficial (blue) and deep (green) layer somata during calcium events, aligned to the time of maximum of all neuropil (**Methods**). Lines represent the hierarchical bootstrap estimates of the mean value for each time bin. Here and elsewhere, shading around the mean is one standard deviation of the bootstrap distribution at each time bin. Response curves are compared for each time bin: in the horizontal bars above the plot, black marks time bins where p<0.05 and gray marks time bins where p>0.05. (**F**) Time of 50% of maximum activity for superficial and deep layer somata during calcium events. (**G**) Top: Calcium fluorescence from the two example somata in **B**. Bottom: Calcium fluorescence from the neuropil surrounding these somata. Fitted slopes during the 60 seconds following calcium event onset are shown as dashed lines. (**H**) Calcium fluorescence of somata (dark gray) and neuropil (purple) in all layers during calcium events. Each soma is aligned to the time of maximum of the neuropil. (**I**) Distribution of the fitted slopes for cells and neuropil during the 60 seconds following calcium events. (**J**) Scatter plot of fitted slopes and correlation of cells with their neuropil. Top and right histograms are marginal distributions over slopes and correlations, respectively.

A third hallmark feature of SD is its preferential propagation through superficial cortical layers (Stafstrom, 2020; Zakharov et al., 2019). To investigate this, we implanted a micro-prism in the cortex, enabling simultaneous imaging across all cortical layers (**Figure 4C, Video S3**). We found that the calcium event first spread in superficial layers 2/3 and 4, and only later reached deeper layers 5 and 6 (**Figure 4D**). Calcium event responses in superficial layers preceded those in deep layers by 6.9 ± 1.3 s (mean ± std) (**Figure 4E-F**). While both somata and neuropil showed a sharp calcium fluorescence increase at the wavefront, calcium decay kinetics were more heterogeneous. Calcium fluorescence in the neuropil gradually returned to baseline over the course of about 100 s. Most somata followed this time course, but a subset of them exhibited a progressive increase in fluorescence (**Figure 4G**), likely reflecting a slow accumulation of somatic calcium. This resulted in a population average fluorescence decrease that was significantly faster in the neuropil than in the somata (**Figure 4H-I**). The somata with increasing post-calcium event fluorescence made up about 35% of the total somatic population (**Figure 4J**). This heterogeneous response pattern is a characteristic of SD, and is hypothesized to be linked to the variable capacity of individual neurons to recover from SD-induced oxidative stress (Enger et al., 2015). Overall, these results strongly support the hypothesis that the ECS-triggered calcium events are indeed SD.

### Calcium events, not ECS, drive Fos expression

If those calcium events are SD, then they should cause neuronal plasticity (Faraguna et al., 2010; Li et al., 2012). To begin testing this hypothesis, we quantified the ECS-driven expression of the Fos protein, the product of an immediate early gene that is a marker of functional plasticity in cortex and hippocampus (Mahringer et al., 2022, 2019). We again performed widefield calcium imaging of ECS responses and allowed one hour of recovery post-ECS for Fos expression (**Methods**). Brain tissue was then processed for Fos immunohistochemistry. We found that ECS drove strong Fos expression in a variety of brain areas, including the dorsal cortex (**Figure 5A**). To determine whether the direct electrical response to the ECS or the subsequent calcium event drove cortical Fos expression, we again took advantage of the occurrence of unilateral calcium events (**Figure 5B**). This allowed us to directly compare Fos expression between hemispheres that exhibited a calcium event and those that did not. Fos was also upregulated in subcortical structures (e.g., amygdala, ventromedial hypothalamus) and in cortical regions outside our widefield imaging field of view (e.g., dentate gyrus, piriform cortex); however, we remain agnostic as to whether the SD invaded these areas. Notably, within our recorded cortical regions, hemispheres without a calcium event exhibited Fos levels that were statistically indistinguishable from sham-treated controls. Instead, the ECS-driven increase in Fos expression in dorsal cortex was fully accounted for by the occurrence of a calcium event (**Figure 5C**). This supports the hypothesis that calcium events, which are likely SD, are the primary driver of neuronal plasticity following ECS.

**Figure 5.**
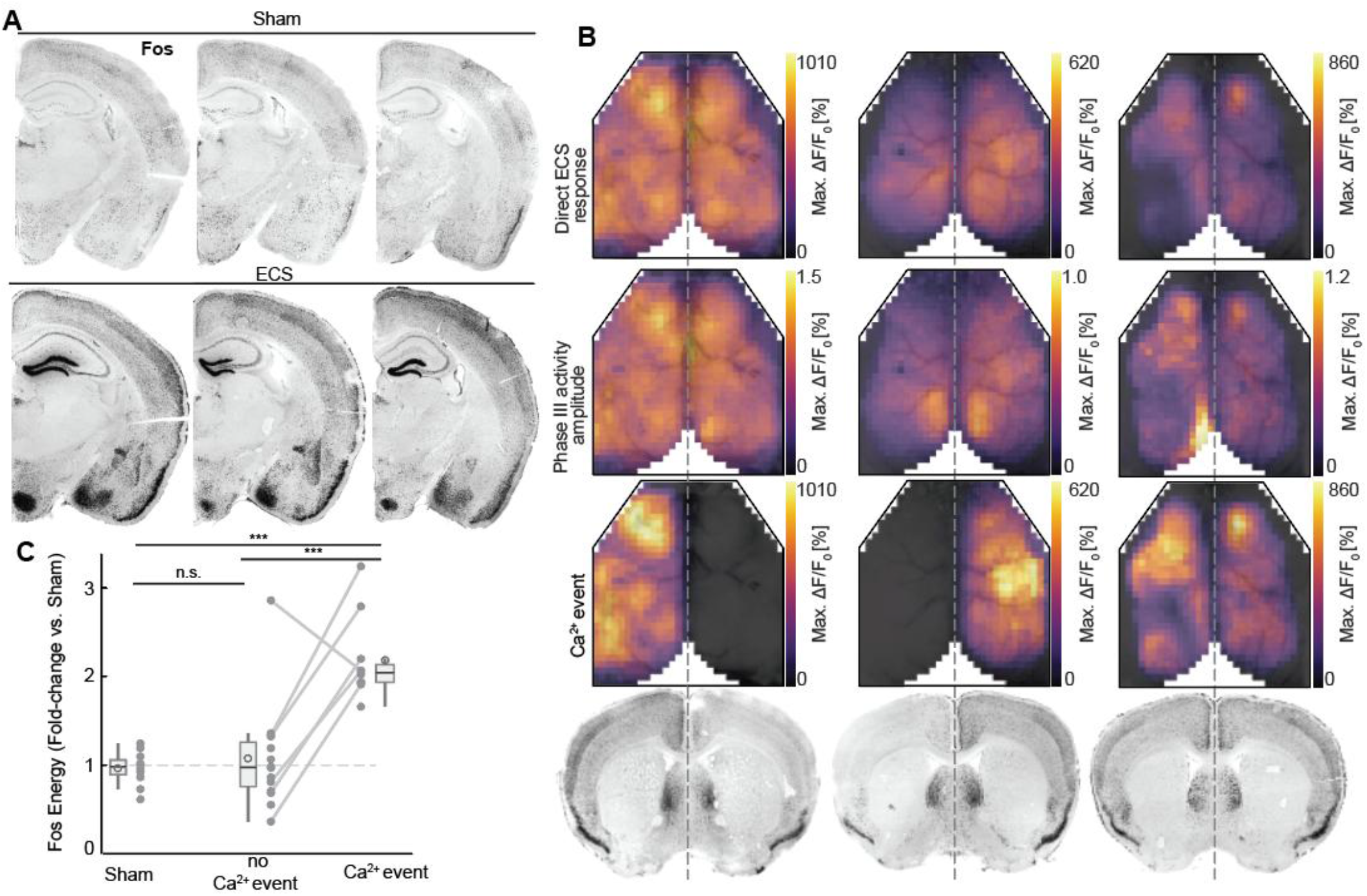
The ECS-triggered increase in Fos expression in dorsal cortex is driven by the calcium event. (**A**) Example of coronal brain sections stained for Fos from three sham ECS (top) and three ECS-treated (bottom) mice. (**B**) Maximum calcium fluorescence during direct ECS response (first row), the amplitude of the ECS-induced phase III activity (second row), and the peak fluorescence to illustrate the presence or absence of a calcium event (third row). From left to right, data from three example mice that had unilateral left, unilateral right, and bilateral calcium events, respectively. The bottom row shows corresponding coronal sections from the same brains, stained for Fos expression. (**C**) Quantification of the Fos throughout the entire cortex, expressed as a fold change for an entire hemisphere normalized to sham mice (see **Methods**). Paired hemispheres (experiments in which ECS triggered a bilateral direct ECS response and phase III activity followed by a unilateral calcium event) are linked by a line. Thus, the calcium event, not the direct ECS response or the phase III activity, is the correlate of Fos expression.

### The occurrence of calcium events correlates with EEG metrics predicting positive ECT therapeutic outcomes

Given that calcium events accounted entirely for the expression of Fos, we asked whether their occurrence might also be associated with known predictors of the therapeutic outcome of ECT. If this were the case, we would expect EEG responses to ECS that correlate with positive therapeutic outcomes to also correlate with the occurrence of calcium events. While EEG responses are only partially predictive of clinical improvement following ECT (Krystal and Weiner, 1994), there are several EEG metrics that correlate with positive therapeutic outcomes. For the phase III activity, an increased amplitude (Perera et al., 2004), a longer duration (Gillving et al., 2024), a higher post ictal suppression index (Kimball et al., 2009; Nobler et al., 1993), and higher interhemispheric coherence (Grözinger et al., 2013), for example, are indicators of a favorable patient response to ECT (Francis-Taylor et al., 2020).

To address this, we investigated whether these EEG metrics correlated with the occurrence of a calcium event, by comparing EEG data from recordings without and with calcium events (**Figure 6A-B**). We found that the mid-phase III activity amplitude was significantly higher in hemispheres where a calcium event occurred compared to the unaffected contralateral hemisphere or to ECS where neither hemisphere showed a calcium event (**Figure 6C**). Both the phase III activity duration and postictal suppression index (**Figure 6D-E**) were increased in ECS triggering a calcium event, but we found no evidence that the laterality of the calcium event (ipsi-vs. contralateral) influenced these two metrics. This absence of hemisphere specificity would be consistent with the observation of a tight coupling of onsets and offsets of phase III activity across hemispheres (Brumback and Staton, 1982). In the case of inter-hemispheric coherence of the EEG signals, we found no evidence of an influence of the calcium event (**Figure 6F**). These results led us to speculate that calcium events may serve as a central mechanistic driver of the therapeutic efficacy of ECT.

**Figure 6.**
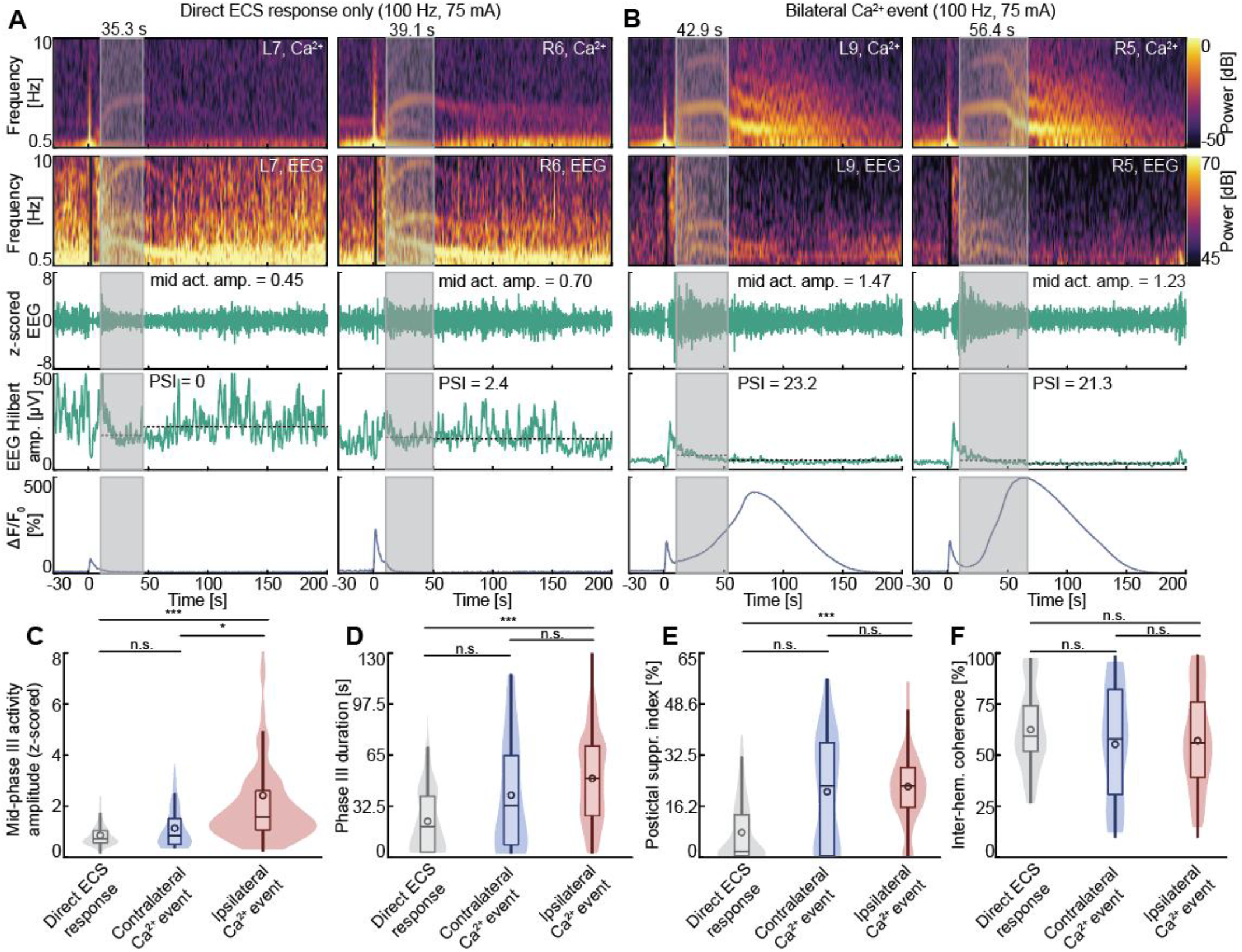
Calcium events are associated with ictal EEG metrics that predict clinical ECT efficacy. (**A**) Example of an ECS session triggering a direct ECS response followed by phase III activity. Data are shown for two example electrodes. A gray transparent square indicates the duration of the phase III activity. First row: Calcium imaging spectrogram of the ROI surrounding the electrode. Second row: EEG signal spectrogram of the electrode. Third row: z-scored EEG signal in the 1-5 Hz band to visualize the phase III activity amplitude. Fourth row: EEG Hilbert amplitude in the 1-30 Hz band, with dashed lines indicating the mean amplitudes during and after the phase III activity used to compute the postictal suppression index (PSI). Fifth row: Calcium imaging fluorescence, low-passed below 0.5 Hz. (**B**) As in **A**, with identical ECS stimulation parameters, but for an ECS session triggering a direct ECS response followed by a bilateral calcium event. (**C**) Mid-phase III activity amplitude for each electrode during ECS sessions triggering only direct ECS response (gray), ECS sessions triggering a unilateral calcium event over the contralateral hemisphere (blue) and ECS sessions triggering a calcium event on the ipsilateral hemisphere (red, either unilateral or bilateral calcium event). See **Methods** for the details on the ictal EEG metrics definitions. (**D**) As in **C**, but for the phase III activity duration. (**E**) As in **C**, but for the postictal suppression index. (**F**) As in **C**, but for the inter-hemispheric coherence of the EEG signal.

## DISCUSSION

Here, we sought to characterize the neuronal response to ECS, focusing on a calcium event that shows hallmark features of an SD. We demonstrate that these events correlate with prognostic EEG markers of ECT treatment outcomes and induce the expression of Fos, a key indicator of neuronal plasticity.

### Limitations

There are three key limitations that constrain the interpretation of our results, and that will need to be addressed in future research:

1. We have no direct evidence that patients also have ECT-induced SD. Patients are typically monitored with two-channel EEG to determine whether ECT has induced a phase III activity (Francis-Taylor et al., 2020), but these measurements are currently not sufficient to detect SD. One simple way to modify the current clinical routine measurements would be to perform direct current (DC) EEG recordings with a higher channel count. SD results in characteristic DC shifts in the EEG response (Dreier et al., 2017; Drenckhahn et al., 2012). While detecting stochastically occurring SD signatures in EEG recordings of awake subjects is difficult (Hofmeijer et al., 2018; Lauritzen et al., 2011), this would likely be facilitated in the context of ECT by the known timing of the triggering stimulus and the elimination of movement artifacts via anesthesia and myorelaxation. Alternatively, given that the SD triggers a strong hemodynamic change (**Figure S5**) (Hadjikhani et al., 2001), blood-oxygenation level dependent imaging techniques might be able to detect an SD. There is already evidence of hemodynamic changes induced by ECT in patients (Rosenthal et al., 2025).
2. Translation may be further hampered by systematic differences in the threshold for triggering SD. Differences in head geometry and stimulation current amplitude between mice and humans could shift the thresholds for both phase III activity and SD. Moreover, the mechanical stress of cranial window surgery may itself altered the susceptibility of cortex to ECS triggered SD in our experiments.
3. Our conclusion that the SD is the primary driver of plasticity is based on its role in driving Fos expression, an early marker of cortical plasticity (Mahringer et al., 2022, 2019). We cannot rule out the possibility that the seizure induced by ECS drives Fos-independent plasticity pathways that also contribute to therapeutic outcomes. Like SD, seizures are neuronal events characterized by supra-physiological excitability that trigger neuronal plasticity (Scharfman, 2002). While knocking out Fos reduces seizure-induced plasticity (Watanabe et al., 1996), other signaling cascades such as mTOR (Kim et al., 2022), BDNF (Ramnauth et al., 2022) and the immediate early gene *Arc* (Larsen et al., 2005) could all be influenced by ECS-induced seizures. One way to explore the contribution of these plasticity pathways to treatment effects will be to map the gene expression changes following ECT more broadly.

### Does ECT drive SD in humans?

In addition to the results presented in this paper and that of (Rosenthal et al., 2025), there are three additional lines of evidence that support the hypothesis that ECT triggers SD in patients:

1. A significant proportion of ECT patients suffer from a migraine-like headache immediately after the treatment (Mulder and Grootens, 2020). Migraines are thought to be caused by an SD-driven activation of Panx1 channels (Karatas et al., 2013).
2. Confusion and cognitive side effects, with a particular focus on memory and hippocampus-dependent processes, are frequently reported following ECT (Impastato and Almansi, 1942; Porter et al., 2020). In mouse models of epilepsy, postictal ambulation has been shown to be driven by hippocampal SD, rather than by the seizure itself (Mitlasóczki et al., 2025).
3. SD causes osmotic changes (Mazel et al., 2002) that could explain the structural imaging changes observed after a single ECT session (Takamiya et al., 2025; Verdijk et al., 2026), which are particularly pronounced in the hippocampus (Bracht et al., 2026; Wilkinson et al., 2017).

Our data show that metrics which predict a positive ECT treatment outcome also correlate with the occurrence of SD in dorsal cortex (**Figure 6**). Given that standard ECT protocols optimize for these metrics as a proxy for seizure quality (Krystal et al., 1995; Sackeim et al., 1991), it is possible that clinical practice has inadvertently optimized ECT to trigger SD. If SD accounts for plasticity effects, triggering an SD in a non-seizure context may be sufficient to elicit therapeutic effects. This is supported by the use of ultrasound stimulation which functions as a treatment for major depressive disorder (Oh et al., 2024) and can trigger SD in mice (Estrada et al., 2021). Much like using a landslide to create ripples in a pond, a generalized seizure may be an unnecessarily burdensome way to trigger the SD required for treatment-relevant plasticity effects. Conversely, the accumulation of many calcium transients over prolonged periods of time may be sufficient to drive immediate early gene expression and trigger treatment-relevant plasticity without SD. This is supported by the cell-type specific increase of small calcium transients during rTMS (Gongwer et al., 2026), which may activate signaling pathways triggered by cumulative calcium load that result in downstream immediate early gene expression, such as the CaMKIV-CREB pathway (Gandolfi et al., 2017; Wu et al., 2001). This makes a testable hypothesis that the effect of rTMS depends on the total calcium influx accumulated over the course of a treatment, consistent with the greater efficacy of higher-dosing stimulation protocols (Cole et al., 2022).

### SD as a driver of ECT-related plasticity

The massive intracellular calcium influx during SD (**Figure 2, 4**) is a prime candidate for driving the molecular changes underlying ECT-induced neuronal plasticity. Calcium is the entry point for a variety of plasticity-related signaling pathways that are disrupted in psychiatric conditions (Nanou and Catterall, 2018). While both physiological activity and seizures can trigger these pathways, SD stands out due to the sheer magnitude of its calcium surge, driven in part by a near-total release of endoplasmic reticulum stores (Stern et al., 2024). Elevated calcium levels drive the expression of key plasticity markers. As such, SD is known to induce Fos in non-ECS contexts (Ingvardsen et al., 1997; Iqbal Chowdhury et al., 2003) and causes differential expression of a host of plasticity-related genes (Dell’Orco et al., 2023). One of the many downstream effectors of these changes is increased BDNF expression (Faraguna et al., 2010). As the primary ligand for the TrkB receptor, BDNF activates a pathway that is a well-established target for both antidepressant (Casarotto et al., 2021) and antipsychotic drugs (Pandya et al., 2013).

Our observation that ECS drives Fos expression in the dorsal cortex following SD contrasts with the reports that ECS fails to increase, or even downregulates, cortical Fos expression following repeated ECS (Winston et al., 1990). There are two possible explanations for this. Because SD occurs stochastically (**Figure 3**), experiments that do not monitor for it cannot attribute the presence or absence of Fos to ECS itself – an ECS without SD results in cortical Fos levels not significantly different from sham ECS (**Figure 5C**). Alternatively, repeated ECS may engage a meta-plasticity mechanism, in which successive inductions progressively dampen Fos expression, for instance through histone deacetylation (Park et al., 2014).

While we cannot fully exclude the possibility of a secondary event that could modulate Fos expression, such as a unilateral seizure restricted to the SD hemisphere, we monitored calcium activity in a subset of mice (11/17) until shortly before euthanasia and did not see any seizure-like calcium activity. We also did not observe behavioral signatures of a seizure in any of the mice. But even if secondary events did occur, the first unilateral event we observed was the SD.

### Origin of phase III activity

In the ECT literature, the phase III activity is often referred to as a seizure. We suspect that the key difference to an epileptic seizure is in the origin of phase III activity and in how the cortex participates in it. An initial strong input to thalamus, either directly from the ECS or indirectly via ECS-driven excitation of cortex, can trigger a 2-4 Hz oscillation (Bal et al., 2000). A central thalamic source is consistent with the observation that phase III activity peaks have zero phase lag across the human cortex (Brumback and Staton, 1982; Huels et al., 2023). At the same time, ECS stochastically triggers SDs. As these move across cortex they silence cortical activity, and what remains in is the oscillatory drive from thalamus. Consistent with this interpretation is the fact that phase III activity can co-exist with SD (**Figure 6B**). This would explain why, following unilateral SD, we found no hemisphere specificity in the postictal EEG suppression (**Figure 6E**). Unlike intracortical LFP, which are thought to reflect local cortical activity (Katzner et al., 2009) and are silenced by SD (Lindquist and Shuttleworth, 2017), the EEG signal we recorded here is likely dominated by the bilateral thalamic drive.

Interestingly, a combination of this phase III activity and SD-induced plasticity could result in a lasting, selective strengthening of thalamocortical synapses. It has been speculated that such an increase is the mechanism that underlies the therapeutic efficacy of ECT (Wei et al., 2020). Reduced coupling between thalamus and frontal areas of cortex is characteristic of schizophrenia (Giraldo-Chica and Woodward, 2017; Vinogradov et al., 2023), and assuming this decoupling is directly related to the etiology of the disease (Keller and Sterzer, 2024), this mechanism of action could explain both the efficacy of ECT in treating psychosis and why patients with psychotic depression have more favorable ECT treatment outcomes than those with nonpsychotic depression (Petrides et al., 2001).

### Future directions

Unresolved is the question of how to reconcile the non-specific nature of SD with the cell-type-specific effects that characterize rTMS (Gongwer et al., 2026), antidepressant (Shao et al., 2025; Vesuna et al., 2020) and antipsychotic drugs (Heindorf and Keller, 2024; Lupori et al., 2026). Several lines of evidence suggest that the effects of SD may also be cell-type-specific. First, SD is easier to trigger in superficial than in deeper layers of cortex (Leao and Morison, 1945), and affects superficial somata for a significantly longer duration (**Figure 4**). This aligns with the evidence that schizophrenia is associated with a number of plasticity-related genes whose expression is selectively enriched in upper cortical layers (Batiuk et al., 2022). Second, the large spectrum of time courses of calcium responses to SD (**Figure 4**) could be explained by cell-type-intrinsic mechanisms, such as differential NMDA receptor expression (Buchanan et al., 2012). Testing this hypothesis will require cell-type-specific chronic calcium imaging following ECS.

The immediate clinical challenge stemming from our findings will be to determine whether the role of SD is therapeutically beneficial or potentially deleterious. Optimizing ECT involves balancing the likelihood of positive therapeutic outcomes while minimizing adverse cognitive effects, most notably retrograde amnesia and impairments in autobiographic memory (Porter et al., 2020; Sackeim et al., 2007). For example, there is an optimal window for the stimulation charge: while a minimum stimulation charge is required for therapeutic benefit, a stimulation charge significantly exceeding the phase III activity threshold is linked to increased cognitive side effects (Delva et al., 2000; Sackeim et al., 1987). This could be explained if a higher stimulation charge is more likely to trigger SDs in brain regions more distant from the stimulation electrodes.

For example, it is conceivable that an SD in frontal cortex is therapeutically beneficial while an SD in the hippocampal-entorhinal system is responsible for retrograde amnesia. If ECS does trigger SD in the hippocampal-entorhinal system, this might open up potential for pharmacological interventions to a local prevention of an SD here, e.g. with NMDA receptor antagonists that preferentially affect hippocampus (Mutel et al., 1998). By restricting SD to frontal neocortex, it may be possible to prevent retrograde amnesia without compromising the cortical plasticity effects required for therapeutic outcomes.

### Relevance

Our findings suggest that SD, a currently unmonitored neuronal response to ECS, appears to be the primary driver of ECT-induced neuroplasticity. Ultimately, we propose that monitoring and facilitating SD during ECT may unlock the possibility to transform psychiatry’s most potent intervention into a targeted and reliable therapy with minimal side effects.

### Ideas and speculation

Based on our results, we propose that SD may be a more relevant biomarker than the phase III activity to explain the therapeutic effects of ECT.

## ACKNOWLEDGEMENTS

We thank all the members of the Keller lab for discussion and support, Tingjia Lu and FMI vector core for AAV production, Jennifer Gröli for animal husbandry, and Felix Widmer for inspiration. Our work was supported by a team of core facilities at the FMI. The project has received funding from the Swiss National Science Foundation (GBK), the Novartis Research Foundation (GBK), EMBO postdoctoral fellowship ALTF 844-2022 (LL), and the European Research Council (ERC) under the European Union’s Horizon 2020 research and innovation programme (grant agreement No 865617) (GBK).

## AUTHOR CONTRIBUTIONS

HL designed the study, performed the widefield calcium experiments and most data analysis. LS, LL, and HL performed the IEG experiments. HL and Ed. S performed the concurrent EEG and widefield calcium imaging experiments. LL, TK and HL performed the two-photon imaging experiments. SU, El. S, GD, and AB performed the ECT treatment in patients and collected the data. All authors wrote the manuscript.

## METHODS

### Mice

A total of 66 mice (27 male, 39 female), 7 – 22 weeks old at the start of the experiment, were used. Most mice were C57BL/6 wild-type, except for transgenic animals used for two-photon prism imaging experiments. In this case, transgene expression was unrelated to the measurements reported here and did not affect experimental readouts. Mice used for two-photon prism imaging were PlexinD1-CreERT2 animals in which ChrimsonR expression had been induced in the retrosplenial cortex (via tamoxifen food). No optogenetic experiment was performed here, and this co-expression does not affect GCaMP stimulation. See **Table S2** for details of mouse inclusion for the different experiments. Mice were group-housed in a vivarium (light/dark cycle: 12/12 hours), and experiments carried out in light cycle. In female mice, we did not time experiments to a specific phase of the estrous cycle. All animal procedures were approved by and carried out in accordance with the guidelines of the Veterinary Department of the Canton of Basel-Stadt, Switzerland.

### Surgery

For all surgical procedures, mice were anesthetized with a mixture of fentanyl (0.05 mg/kg; Actavis), midazolam (5.0 mg/kg; Dormicum, Roche), and medetomidine (0.5 mg/kg; Domitor, Orion) injected intraperitoneally. Analgesics were applied perioperatively (2% lidocaine gel, meloxicam 5 mg/kg) and postoperatively (3.25 mg/kg; Ethiqa XR). Eyes were covered with ophthalmic gel (Virbac Schweiz AG). For widefield imaging, we implanted crystal skull (Kim et al., 2016) or thinned skull (Yang et al., 2010) cranial windows. Briefly, the perimeter of the skull plate overlying the dorsal cortex was thinned using a dental drill until it could be removed with forceps (crystal skull) or only thinned in its entirety until transparent but not removed (thinned skull). To protect the brain, a glass or acrylic polymer window was then implanted. For two-photon imaging, a cranial window was implanted over primary visual cortex (V1) as previously described (Keller et al., 2012; Leinweber et al., 2014). Briefly, a 4 mm craniotomy was made over the right V1, centered 2.5 mm lateral and 0.5 mm anterior to lambda. The exposed cortex was sealed with a 3 mm or 4 mm circular glass coverslip. For prism imaging, a 0.7 mm glass prism (Tower Optical Corporation) was attached to the coverslip with UV-curable glue (Norland Optical Adhesive), and a small incision of cortical tissue was made to allow for the insertion of the prism into the cortex. Cranial windows were sealed using gel superglue (Ultra Gel, Pattex), and, for widefield windows, cyanoacrylate adhesive (Vetbond, 3M) (Kauvar et al., 2020). The remaining exposed surface of the skull was covered with cyanoacrylate adhesive, and a titanium head bar was fixed to the skull using dental cement (Paladur, Heraeus Kulzer). After surgery, anesthesia was antagonized by a mixture of flumazenil (0.5 mg/kg; Anexate, Roche) and atipamezole (2.5 mg/kg; Antisedan, Orion Pharma) injected intraperitoneally.

### Virus injections

For widefield experiments, we deposited an AAV vector with PHP.eB capsid (Chan et al., 2017) retro-orbitally (4 μl per eye of at least 10^14^ GC/ml) to drive expression throughout cortex of GCaMP6s (AAV-PHP.eB-Ef1α-GCaMP6s-WPRE) or EGFP (AAV-PHP.eB-Ef1α-EGFP-WPRE). For two-photon experiments, the AAV was injected locally in the primary visual cortex using a pneumatic injector (IM-300 Microinjector, Narishige) to drive expression of GCaMP6f (AAV2/9-Ef1α-GCaMP6f-WPRE, 100-250 nL of at least 10^14^ GC/ml). Imaging commenced at the earliest 3 weeks after AAV injection.

### Electroconvulsive stimulation (ECS) in mice

For all ECS experiments, mice were anesthetized with isoflurane (2%) using 95% O_2_ as a gas carrier (1 L/min). For myorelaxation we injected succinylcholine chloride (1 mg/kg; Sigma-Aldrich) intraperitoneally. Mice were head-fixed, and ECS was delivered through auricular electrodes (Ugo Basile model #57802) using a pulse generator (Ugo Basile model #57800-001). A conductive salt-free electrolyte gel (Spectra 360 gel, Parker Laboratories) was applied to the electrodes to improve conductivity. Stimulation consisted of a 1 s long pulse train with a frequency of between 25 and 100 Hz, and a current amplitude of between 15 and 75 mA. Pulse width was 0.5 ms, which resulted in a total charge range of between 0.187 and 3.750 mC. These parameters reliably evoke tonic-clonic convulsions in mice (Hagen et al., 2015; van Buel et al., 2017). The distribution of ECS parameters and the specific parameters used for each single ECS example can be found in **Table S3**.

For comparison, patient ECT parameters used here were typically as follows: train duration up to 8 seconds, pulse frequency of between 40-70 Hz, current amplitude of between 890-920 mA, pulse width of between 0.5-0.75 ms, charge maximum 1000 mC. Thus, stimulus duration, current amplitude and resulting total charge are adapted to the mouse, but frequency/pulse-width are within clinical range (Peterchev et al., 2010). The list of ECT parameters used here can be found in **Table S4**. The stimulation polarity was constant for mice ECS, whereas bidirectional pulse train (alternating polarity) was used for patient ECT. For sham ECS experiments, we used an identical procedure with no current delivery. To limit adaptation to repeated stimulation, each mouse received at most 6 ECS sessions administered at intervals of 48 hours. Given this constraint, we sampled stimulation parameters space more densely at higher frequencies and current amplitudes where calcium events were more likely. As a result, the number of stimulations per parameter combination was not balanced (**Figure 3E**). However, stimulation parameters were randomized within and across mice to minimize order effects.

Of note, some prior studies have used epicranial electrodes for mouse ECS (Rosenthal et al., 2025). This approach builds on intracranial work in awake rats and uses mismatched stimulation parameters, which limits its translational validity (Theilmann et al., 2014). In contrast, the auricular ECS used here matches the non-invasiveness of ECT in patients and triggers patient-like phase III activity (**Figure 1**). The key difference is that epicranial stimulation bypasses the skin and connective tissue. In the rat brain, electric fields generated by skull stimulation are almost an order of magnitude larger than those generated by skin stimulation and resemble those generated by direct stimulation of the brain surface (Asan et al., 2019).

### Electroconvulsive therapy (ECT) in humans

The retrospective treatment data of 26 random patients (8 male, 18 female, all > 18 years old), who underwent ECT at the Zentrum für Affektive, Stress- und Schlafstörungen (ZASS), University Psychiatric Clinics (UPK), Basel due to a psychiatric indication were included. Patients signed a general consent form (approved by the local ethics committee). All patient procedures were approved by and carried out in accordance with the Ethikkommission Nordwest- und Zentralschweiz (EKNZ). One randomly chosen treatment within the first half of the ECT acute treatment series (i.e., < 7th treatment) per person was considered for analysis. Restimulation treatment data were excluded. Medical indication for ECT, patient information, treatment preparation and anesthesia followed standardized clinical protocols according to current standards of care. ECT was conducted with bifrontal stimulation using the Thymatron® System IV device (Somatics, LLC, Lake Bluff, IL, USA) and disposable adherent stimulus electrodes (EPAD, Thymapad®), as previously described (American Psychiatric Association, 2024), following Kellner’s ECT guidelines (Kellner, 2018). Briefly, patients were anesthetized with either propofol or etomidate (Braun Medical AG) and myorelaxed with succinylcholine chloride (Amino AG). Skin surface was cleansed using an alcohol-containing solvent and abrasive gel. A conductive and adhesive liquid (Signaspray®, Parker Laboratories) was then applied to the skin. EEG was recorded at 200 Hz using bilateral frontomastoid placement and filtered using a band-pass with 3 dB points at 2 and 25 Hz. We excluded the first two seconds of recording following the stimulation to remove a capacitive coupling artifact. ECT was performed about 4 min after the injection of etomidate and 5 min after the injection of propofol, using the ‘double dose’ stimulation protocol. ECT sessions included in the analysis were performed by multiple clinicians with standardized preparation and stimulation procedures. Data were measured using the Thymatron during each session, including stimulation parameters and EEG-derived seizure quality parameters. In addition, we had access to formalized clinical documentation listing the administered medication and manually assessed seizure duration. Digitalization of stored ECT data for the current analysis was performed after December 2020 using GENET-GPD (Freundlieb et al., 2023). The digitalized treatment data were derived from the GENET-GPD database. Patient data were stored in the UPK patient case files in compliance with accepted data protection and record-keeping regulations.

### Electroencephalography (EEG) in mice

Mice were anesthetized as described above, and the glass or polymer window was removed. An EEG multielectrode array with 30 recording sites (EEG mouse-30-A-10-H32, Neuronexus) was used to record neural activity across the surface of the dorsal cortex. Signals were amplified using a 32-channel headstage (RHD 32ch, Intan Technologies) and digitized at 30 kHz using an Open Ephys acquisition board (Siegle et al., 2017). During ECS delivery, the EEG array was physically disconnected by an electronically triggered relay to prevent electrical arcing between electrodes. EEG data were downsampled to 1 kHz and band-pass filtered between 0.5 and 10 Hz. We excluded the first two seconds of recording following EEG reconnection to remove a capacitive coupling artifact (**Figure S1**). As the concurrent widefield calcium signal provided an unambiguous polarity reference, the polarity of the EEG signal was such that positive voltage deflections corresponded to increases in fluorescence.

### Widefield imaging

Widefield imaging experiments were conducted on a custom-built macroscope with objectives mounted face-to-face (Nikon 85 mm/f1.8 sample side, Nikon 50 mm/f1.4 sensor side). A 470 nm LED (Thorlabs) powered by a custom-built LED driver was used to excite fluorescent indicators through an excitation filter (SP490, Thorlabs) reflected off of a dichroic mirror (LP490, Thorlabs) placed parfocal to the objectives. The fluorescence was collected through a 525/50 emission filter on a sCMOS camera (PCO edge 4.2). LED illumination was adjusted with a collimator (Thorlabs SM2F32-A) to homogeneously illuminate cortical surface through the cranial window. Images were acquired at 100 Hz frame rate. Raw images were cropped on the sensor and data were stored to disk with custom-written LabVIEW (National Instruments) software, resulting in an effective pixel size of 60 μm^2^ at a resolution of 1108 by 1220 pixels (1.35 MP).

### Two-photon imaging

Two-photon calcium imaging was performed using custom-built microscopes (Leinweber et al., 2014). The illumination source was a tunable femtosecond laser (Insight, Spectra Physics or Chameleon, Coherent) tuned to 930 nm. Emission light was band-pass filtered using a 525/50 filter and detected using a GaAsP photomultiplier (H7422, Hamamatsu). Photomultiplier signals were amplified (DHPCA-100, Femto), digitized (NI5772, National Instruments) at 800 MHz, and band-pass filtered at 80 MHz using a digital Fourier-transform filter implemented in custom-written software on an FPGA (NI5772, National Instruments). The scanning system of the microscopes was based on a 12 kHz resonant scanner (Cambridge Technology). Images were acquired at a resolution of 750 by 400 pixels (60 Hz frame rate) with a 16x, 0.8 NA water immersion objective (Nikon). We imaged layer 2/3 at a depth of 100-250 μm with a field of view of 375 by 300 µm. For prism imaging, to enable imaging of as much cortical depth as possible, the field of view was expanded to 270 by 960 µm, at a resolution of 750 by 800 pixels. This resulted in an effective frame rate of 30 Hz. Laminar boundaries were estimated from the density of GCaMP6-expressing cells, which typically peaks in layer 2/3 and upper layer 5 (Andermann et al., 2013).

### Image processing

For analysis of chronic widefield macroscope imaging, raw movie data were manually registered across days by aligning subsequent mean projections of the data to the first recorded image sequence. In the raw 1108 by 1220 pixel images, we tiled the dorsal cortex with 818 ROIs each with a fixed location relative to bregma and lambda. This resulted in an average ROI size of 25 by 25 pixels (approximately 230 by 230 μm). Each ROI was also registered against the Allen Mouse Brain Common Coordinate Framework (Wang et al., 2020). In concurrent EEG and widefield recordings, we defined annular ROIs centered on each of the 30 electrode sites. The inner boundary corresponded to the electrode footprint with a 50 μm buffer (approximately 60 pixels in diameter; approximately 550 μm) and was excluded from analysis, while the outer boundary defined a circular region with a diameter of approximately 80 pixels. Two-photon calcium imaging data were processed as previously described (Attinger et al., 2017; Keller et al., 2012). Briefly, raw images were full-frame registered to correct for lateral brain motion. Soma ROIs were manually selected based on mean and maximum fluorescence images. For each soma ROI, a corresponding neuropil ROI of equal pixel area was created as a surrounding annulus, maintaining a 3-pixel exclusion buffer and avoiding overlap with other defined ROIs. No neuropil subtraction to somata activity was done. Activity was calculated as the ΔF/F_0_, where F_0_ was the median fluorescence of the recording before ECS (typically 30 seconds). ECS recordings lasted for 5 min.

### Histology

Mice underwent a single ECS or sham procedure while being monitored for calcium events through widefield imaging and were returned to their home cage for one hour. A subset (11/17) of mice was instead kept head-fixed and imaged for 1 hour to verify that no additional calcium events occurred. Mice were then anesthetized with a mixture of fentanyl (0.05 mg/kg; Actavis), midazolam (5.0 mg/kg; Dormicum, Roche), and medetomidine (0.5 mg/kg; Domitor, Orion), and transcardially perfused with phosphate-buffered saline (PBS) solution and subsequently with ice-cold 4% paraformaldehyde (PFA) in PBS. Brains were extracted and post-fixed overnight in 4% PFA at 4 °C. They were then embedded in 4% agarose and coronally sectioned through the entire anteroposterior extent of the cortex at 50 µm thickness using a vibratome (Leica VT1000S). We collected every fourth section for further processing, resulting in an effective sampling step of 200 µm. Free-floating sections were then processed for Fos immunohistochemistry. Briefly, sections were blocked for 2 h at room temperature in a solution containing 5% donkey serum (Merck Millipore) and 0.5% Triton X-100 in PBS, then incubated overnight at 4 °C with a solution containing anti-Fos antibody (226008, Synaptic Systems, 1:1000), 1% donkey serum and 0.1% Triton X-100 in PBS. Wells were then rinsed 3 times in PBS (10 min each) and incubated with a solution containing donkey polyclonal anti-rabbit antibody (A31573, Invitrogen, Alexa Fluor™ 647 conjugate, 1:300), 1% donkey serum and 0.1% Triton X-100 in PBS for 2 h at room temperature. Following additional 10-minute PBS washes, nuclei were counterstained with DAPI (Sigma-Aldrich, 1:1000) for 10 minutes. Finally, sections were mounted on microscopy slides with a mounting medium (VECTASHIELD, H-1000, Vector Laboratories), sealed with nail polish, and stored at 4 °C. Images were then acquired with an Axioscan Z1 (Zeiss) microscope using an apochromatic 10x/0.5 M27 objective (Zeiss). Fluorophores were excited with 422 nm and 668 nm X-Cite Xylis LED. The emission wavelengths were filtered with a BP 445/50 filter for DAPI and a BP690/50 filter for Alexa 647. Images were collected with a Hamamatsu ORCA flash 4.0V camera at a pixel size of 0.650 µm.

### Tissue registration and quantification

Images were aligned to the Allen Mouse Brain Common Coordinate Framework (CCFv3-2017) as described previously (Lupori et al., 2023). All preprocessing and registration steps, unless stated otherwise, were carried out using custom-written MATLAB (MathWorks) code. For each brain, images were ordered along the posterior-to-anterior direction. Based on a reference mark identifying the right side and/or tissue irregularities, images were mirrored vertically so that all matching hemispheres would be on the same side. Binary masks to restrict quantification to brain tissue were automatically generated from the DAPI images by training a pixel classifier in Ilastik (Berg et al., 2019). Each mask was then visually inspected and manually adjusted based on the Fos images, to remove regions containing tissue damage or staining artifacts. Left-right binary masks defining which hemisphere a pixel belongs to were then generated by manually drawing a segment at the midline of each slice. Using the DAPI images, each slice was matched to a plane in the CCFv3 through a first step of global alignment using QuickNII v2.2 (Puchades et al., 2019) and a second step of local non-rigid transformation with VisuAlign v0.9 (RRID: SCR_017978, VisuAlign). For cell counting, Fos channel images were first pre-processed with a rolling ball filter (50 pixels or 32.5 μm radius) for background removal. Cell detection and counting were then performed on the filtered images with custom scripting inside QuPath v.0.6.0 (Bankhead et al., 2017). Briefly, we used the inbuilt automatic cell detection tool to identify Fos-positive cells using the median fluorescence of the slide as a threshold. A random forest classifier was then trained to remove detection artifacts. For each cell, we calculated mean pixel intensity and x-y coordinates of the centroid. For all regions descending from isocortex, as per the hierarchy defined in the CCFv3, we computed cell density (number of cells/total area in mm2), mean cell intensity, and energy (cell density*mean cell intensity) (Lein et al., 2007; Lupori et al., 2023). For each metric, the value was then normalized by the average of that metric across all sham-treated mice.

### Data analysis

All data analysis was performed with custom-written MATLAB (MathWorks) code.

When visualizing data with a box plot, the central box indicates the interquartile range (IQR), spanning from the first (Q_1_) to the third (Q_3_) quartiles. Whiskers extend to data points not considered outliers. Outliers were defined as data points lying outside of the Q_1_ – 1.5 IQR to Q_3_ + 1.5 IQR range. A central horizontal line indicates the median and an open circle the mean of the distribution. To represent the underlying probability density, a kernel density estimate was computed using a Parzen-window method with a normally distributed kernel.

In **Figure 1** and **Figure 6**, spectrograms were computed using a 2 s sliding window with 95% overlap of the band-passed signal between 0.5 and 10 Hz. Power spectral density was estimated for each channel or ROI independently and converted to decibel scale. Phase III activity was detected in the 1-5 Hz band by computing the Hilbert envelope of the EEG signal. Activity onset was defined as the first crossing of the 90^th^ percentile of the envelope after ECS. Activity termination was defined as the first time the envelope dropped below a baseline threshold. Patient EEG is interrupted rapidly after the termination of phase III activity. To account for the differences in recording length, the termination threshold was set as the 50^th^ percentile Hilbert envelope for mice and as the 5^th^ percentile for patients. The Hilbert envelope was smoothed with a moving average filter of 10 s for mice and 2 s for patients. For patient EEG, this detection method yielded activity durations not statistically different from those computed by the proprietary Thymatron algorithm (hierarchical bootstrap with resampling, p = 0.268). Power-weighted mean frequencies were computed as the sum of each frequency multiplied by its power, divided by the total power in the 1-5 Hz band. We reported the mean values in a 2 s window centered on the global frequency peak and in the final 2 s of the detected phase III activity. For mouse widefield data, the onset and offset times were computed from the concurrent EEG signal. We excluded frequencies between 7 and 9 Hz, if they exceeded 90 μV; these regions were dilated by 2 s and masked-out from further analysis. We completely excluded EEG electrodes and their associated widefield ROI if the onset/offset values were non-finite or if the total duration of phase III activity was less than 1 s, resulting in the exclusion of 78/480 (16%) of electrodes in our dataset.

In **Figure 2** and **Figure S3**, calcium event speed was estimated through the local fluorescence gradient. Peak latencies were defined as the global maximum occurring 10 to 200 seconds after local wavefront onset. For each ROI, raw fluorescence values were low-passed filtered below 0.5 Hz and a plane (*t* = *S_x_x* + *S_y_y* + *t*_0_) was fitted to the peak latencies of neighbors within a 0.8 mm radius using bisquare weighting. This determined a local slowness vector, the magnitude of which was inverted to determine speed. Speed values deviating more than 3.5 median absolute deviation from the session median were excluded. For two-photon imaging, propagation speed was calculated as the median velocity of calcium events across pairs of somata ROIs separated by at least 200 µm.

In **Figure 2** and **Figure S4**, the period of direct ECS stimulation is defined as the first 5 s post ECS onset, based on the duration reported in **Figure S1**.

In **Figure 2, 3** and **Figure S3**, the onset of a calcium event was defined as the first crossing of 150% of the mean calcium activity before ECS. Only events occurring 10 s after ECS were considered as calcium events. For analysis of time to peak, we excluded ROIs for which there was no clear separation between direct ECS response and calcium event. To determine the presence of a phase III activity, we averaged the calcium traces across hemispheres and used the same algorithm used in **Figure 1** and **Figure 6**. To characterize the probability of calcium event or phase III activity as a function of ECS parameters, we fitted a linear slope with ordinary least squares method. 95% confidence bands for the mean predicted response were computed across the range of predictor variables.

In **Figure 2**, to generate isochrones, we set an ROI-wise threshold to 200% of the maximum ECS baseline activity. The first time at which each ROI crossed this threshold was used to construct a time-of-activation map, which was then filtered with a normal distribution. Contour lines were extracted at 2 second intervals from the onset of calcium event until 50 s post-onset.

In **Figure 4**, cortical layers were separated into supragranular and infragranular, based on empirical distribution of cell densities, which shows a clear gap in cortical layer 4. To compare the activity of somata with their neuropil, we aligned each soma with the peak of its neuropil during calcium events. This alignment ensures that the propagation delay across the field of view is not taken into account. To compare supragranular and infragranular layers, somata were instead aligned to the max of a neuropil trace, averaged across all neuropil ROIs in the entire recording. To quantify the dynamics of GCaMP6f fluorescence following the calcium event, we computed a linear recovery slope with least-square ordinary fit in the 2 to 60 seconds following peak somata signal.

In **Figure 6**, the mid-ictal amplitude was computed as the mean of all phase III activity peaks in a 4 s window centered at the mid-duration of the phase III activity (Krystal et al., 1995). The postictal suppression index was defined as the ratio of the median 1-30 Hz envelope amplitude post-phase III activity over intra-phase III activity. Welch’s periodogram method with a 1 s Hann window and 0.5 s overlap was used to compute the inter-hemispheric coherence between pairs of symmetric electrodes. We reported the values of coherence in the 1 to 5 Hz band.

In **Figure S2**, to illustrate the synchronicity of EEG and calcium signals, peaks in the EEG channel were detected using the Automatic Multiscale Peak Detection algorithm, which is suited to non-stationary signals, such as EEG, without requiring manual parameter tuning or pre-filtering (Scholkmann et al., 2012).

### Statistical analysis

All statistical information for the tests performed in the manuscript is provided in **Table S1**. Analyses accounted for the nested structure of the data consisting of recording units (neurons, EEG channels or ROIs) within subjects (mice or patients) using hierarchical bootstrap (Saravanan et al., 2020). Briefly, we first resampled with replacement data at the subject level and then resampled with replacement a second time at the recording unit level. We then computed the mean of this bootstrap sample and repeated this N times to generate a bootstrap distribution of the mean estimate. For all statistical testing the number of bootstrap samples (N) was 10 000, for plotting bootstrap mean and standard error response curves it was 1000. The bootstrap standard error is the 68% confidence interval (1 SD, 15.8^th^ percentile to 84.2^nd^ percentile) in the bootstrap distribution of the mean.

To account for the false discovery rate in multiple comparisons, we applied an adjustment to p-values (Benjamini and Hochberg, 1995). Briefly, p-values were sorted in ascending order and multiplied by the ratio of the total number of comparisons within a data group (for example, all bootstrap tests for a specific EEG metric) to their sorting rank. The final significance cutoff was determined by finding the largest p-value smaller than its rank-specific threshold, at which point all p-values with a higher rank (i.e., those with smaller p-values) were also rejected.

For linear regression analysis, we computed the F-statistic to test the significance of the model. This test evaluates whether the slope is significantly different from zero by comparing the model degrees of freedom (the number of predictors, here, one) against the degrees of freedom for error (the number of observations minus the parameters for the intercept and slope). The resulting F-statistic represents the ratio of explained variance to unexplained variance, quantifying the portion of the data’s variability accounted for by the linear regression. For multivariate linear regression analysis, we measured the probability of occurrence of a calcium event as a function of stimulation amplitude, frequency, and their two-way interaction. For each of those three parameters, we performed a two-sided t-test against the null hypothesis that this parameter is equal to zero.

### Data and code availability

Software for controlling the two-photon and widefield microscopes and preprocessing of calcium imaging data is available on https://sourceforge.net/projects/iris-scanning/. All mouse data and code necessary to generate the figures of this manuscript will be deposited upon acceptance to zenodo.org. Patient data will be made available upon reasonable request.

---

**Figure S1.**
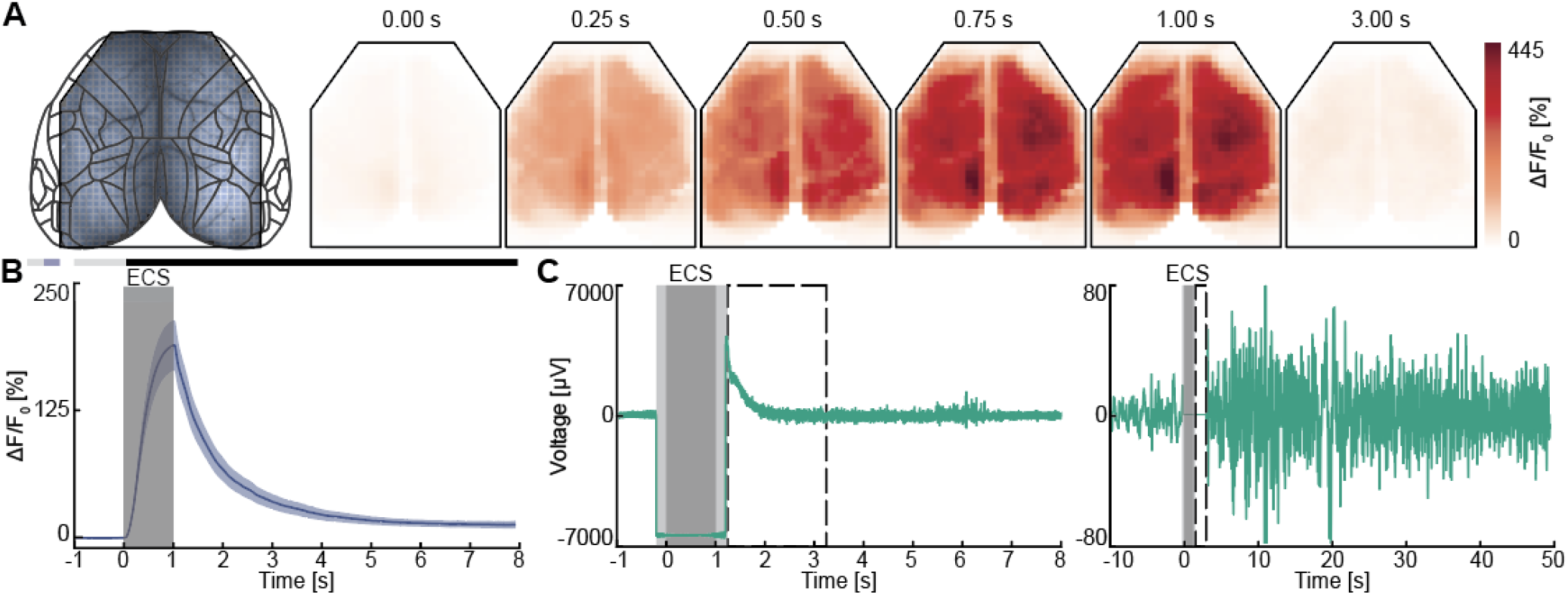
ECS triggers a direct calcium response during the EEG disconnection. Related to Figure 1. (**A**) Left: Widefield recordings (without concurrent EEG) were parcellated into 818 ROIs, each of which was assigned to a cortical region according to the Allen Brain Atlas taxonomy. Right: Direct ECS calcium activity triggered by an example ECS. Each panel shows the average calcium activity over a 10 ms window starting at the time indicated above the panel. (**B**) Direct ECS calcium response estimated using hierarchical bootstrap across all ECS. Response curves are compared against baseline distribution for each time bin: in the horizontal bars above the plot, black marks time bins where p<0.05 and gray marks time bins where p>0.05. Duration of ECS is marked as a dark gray shaded area. (**C**) Left: during ECS (dark gray shaded area), the EEG electrode is temporarily disconnected to prevent arcing (**Methods**), leading to a period in which no EEG data is recorded (light gray shaded area). To eliminate the capacitive discharge artifact upon reconnection, the first 2 s of data are excluded from analysis (dashed box). Right: same signal with a longer time scale, following artifact removal and 0.5–10 Hz band-pass filtering.

**Figure S2.**
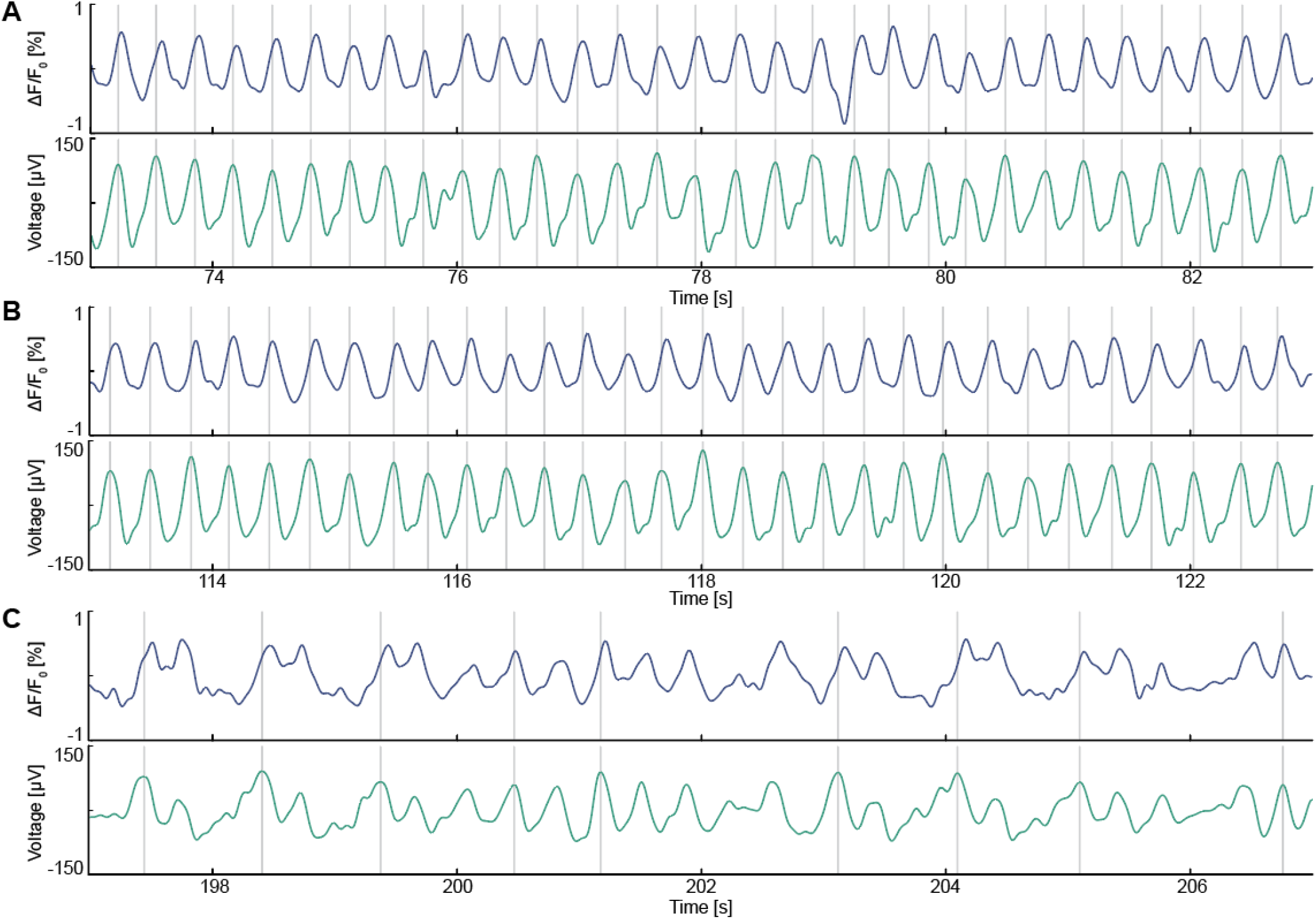
ECS-triggered activity is synchronized between widefield calcium and EEG in mice. Related to Figure 1. (**A**) Activity observed early after ECS in an example mouse widefield calcium imaging ROI (blue) and in the concurrently recorded EEG (green). Activity peaks were detected using Automatic Multiscale Peak Detection on the EEG channel (**Methods**). (**B**) As in **A**, in a period of time 113 s after ECS. (**C**) As in **A**, in a period of time 197 s after ECS.

**Figure S3.**
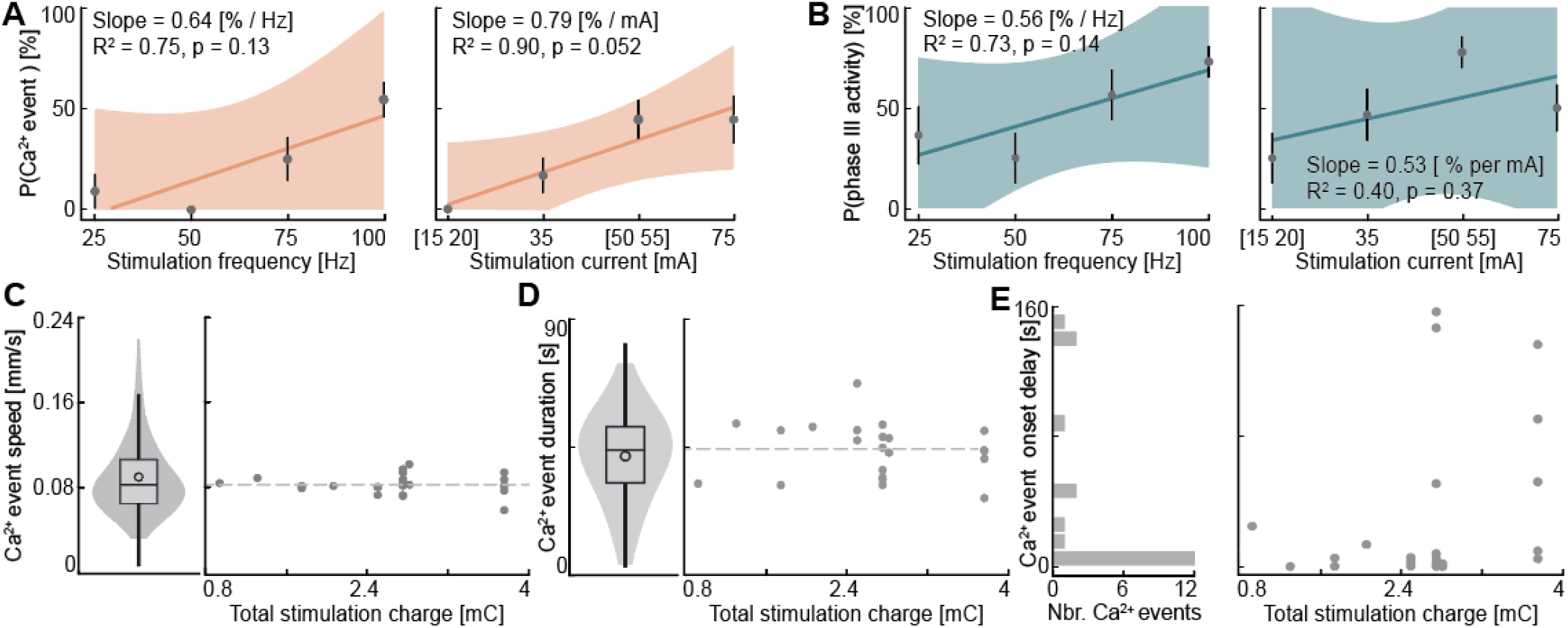
Influence of stimulation parameters on calcium events. Related to Figure 3. (**A**) Probability of triggering a calcium event as a function of stimulation frequency (left) or current (right). The solid line represents the linear fit of the data; the shaded region represents 95% confidence interval on the fit. Error bars on individual data points indicate standard error. (**B**) As in **A**, but for phase III activity. (**C**) Left: Distribution of calcium event speed across triplets of ROIs (see **Methods**). Right: Median calcium event speed as a function of stimulation charge. Dashed line indicates median calcium event speed across all charges. (**D**) Left: Distribution of the duration of calcium events in each widefield ROI (see **Methods**). Right: Median duration of calcium event as a function of stimulation charge. Dashed line indicates median duration across all charges. (**E**) Left: Delay from stimulation to calcium event onset. Right: Delay from stimulation to calcium event onset as a function of stimulation charge.

**Figure S4.**
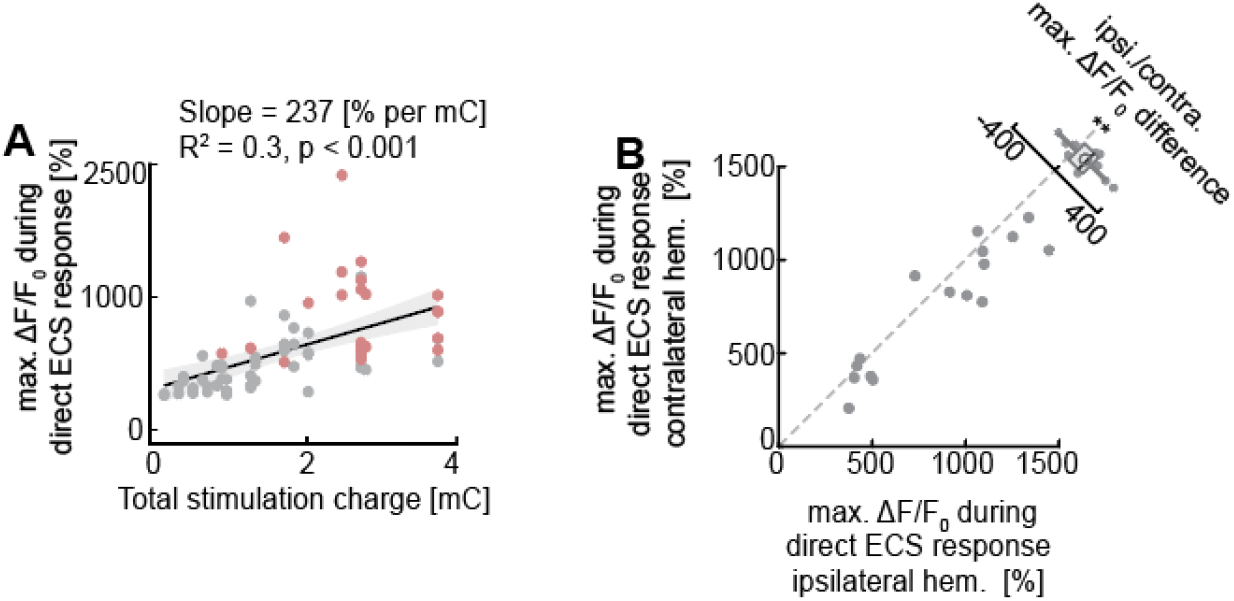
Direct ECS response in calcium imaging predicts the occurrence of calcium events. Related to Figure 3. (**A**) Left: Maximum calcium activity during the direct ECS response as a function of stimulation charge. Red values are measured during ECS triggering a calcium event (either unilateral or bilateral) and gray values from ECS that did not trigger a calcium event. (**B**) Maximum calcium activity in both hemispheres during the direct ECS response, before a unilateral calcium event was triggered.

**Figure S5.**
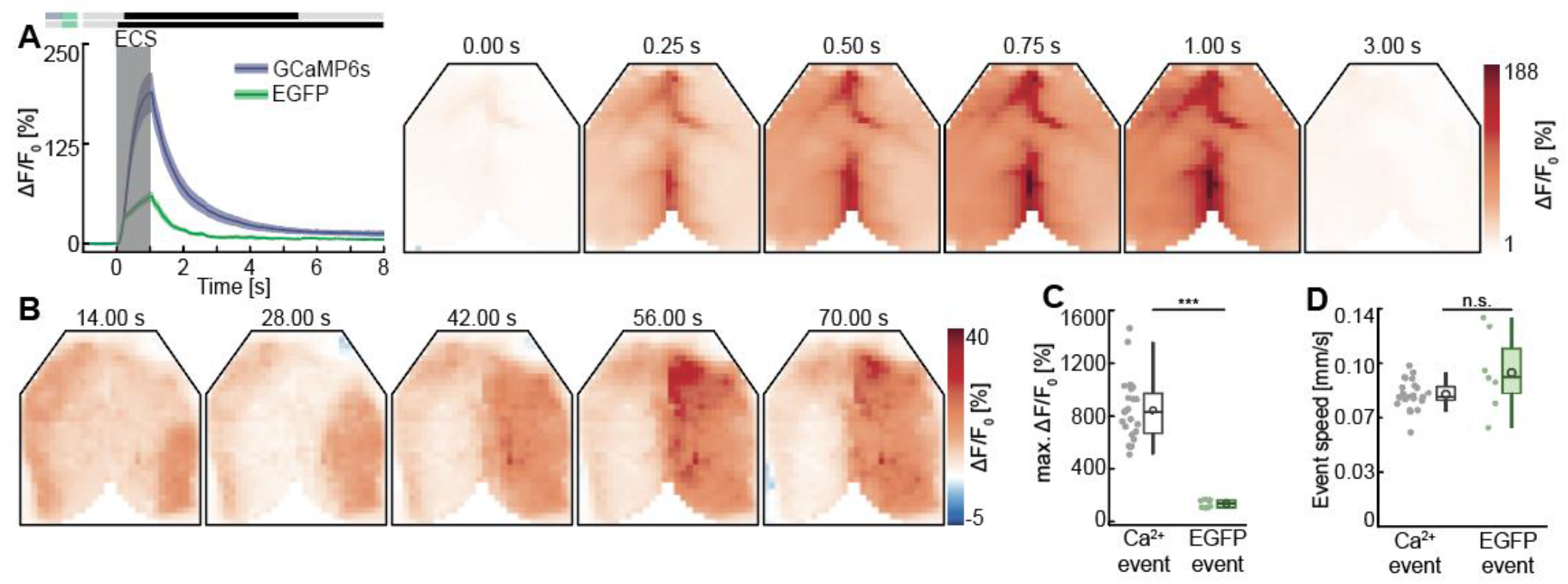
ECS triggers hemodynamic responses. Related to Figure 3. (**A**) Left: Direct ECS fluorescence measured with GCaMP6s (as shown in **Figure S1B**) and EGFP, estimated across the population using hierarchical bootstrap. In the horizontal bars above the plot, black marks time bins where p<0.05 and gray marks time bins where p>0.05. The top bar represents the comparison between GCaMP6s and EGFP fluorescence, while the bottom bar compares the EGFP fluorescence to baseline distribution. Right: Direct EGFP fluorescence evoked by a single example ECS. Each panel shows the average fluorescence over a 10 ms window starting at the time indicated above the panel. (**B**) Low-passed EGFP fluorescence during an example ECS session with unilateral calcium event-like propagation over the right hemisphere. (**C**) Maximum ΔF/F0 during calcium and EGFP events. (**D**) Propagation speed of traveling events (**Methods**). Each data point is the median speed across one event.

**Video S1. Widefield calcium imaging of a calcium event following ECS.**

Examples of ECS triggering a calcium event over the left hemisphere (left), both hemispheres (middle) and right hemisphere (right) in widefield calcium imaging.

**Video S2. Two-photon calcium imaging of a calcium event following ECS.**

Examples of ECS triggering a calcium event while imaging upper L2/3 with two-photon calcium imaging.

**Video S3. Two-photon micro-prism imaging of a calcium event following ECS.**

Examples of ECS triggering a calcium event while imaging all cortical layers through a micro-prism implant (see **Methods**).

## Key Resource Table

| REAGENT OR RESOURCE | SOURCE | IDENTIFIER |
|---|---|---|
| **Bacterial and virus strains** |  |  |
| AAV-PHP.eB-EF1α-GCaMP6s-WPRE (10<sup>14</sup> GC/ml) | FMI vector core | vector.fmi.ch |
| AAV-PHP.eB-EF1α-EGFP-WPRE (10<sup>14</sup> GC/ml) | FMI vector core | vector.fmi.ch |
| AAV2/9-EF1α-GCaMP6f-WPRE (10<sup>14</sup> GC/ml) | FMI vector core | vector.fmi.ch |
| AAV2/9-hSyn-DIO-ChrimsonR-tdTomato (10<sup>14</sup> GC/ml) | FMI vector core | vector.fmi.ch |
| **Chemicals, peptides, and recombinant proteins** |  |  |
| Fentanyl citrate | Actavis | CAS 990-73-8 |
| Midazolam (Dormicum) | Roche | CAS 59467-96-8 |
| Medetomidine (Domitor) | Orion Pharma | CAS 86347-14-0 |
| Ropivacaine | Fresenius Kabi | CAS 132112-35-7 |
| Lidocaine | Bichsel | CAS 137-58-6 |
| Buprenorphine hydrochloride (Ethiqa XR) | Fidelis Pharmaceuticals | CAS 53152-21-9 |
| Ophthalmic gel (Humigel) | Virbac | N/A |
| Flumazenil (Anexate) | Roche | CAS 78755-81-4 |
| Atipamezole (Antisedan) | Orion Pharma | CAS 104054-27-5 |
| N-Butyl-2-cyanoacrylate (Vetbond) | 3M | CAS 6606-65-1 |
| Dental cement (Paladur) | Heraeus Kulzer | CAS 9066-86-8 |
| Spectra 360 electrode gel | Parker Laboratories | N/A |
| Attane Isoflurane | Provet AG | CAS 26675-46-7 |
| Succinylcholine chloride | Sigma-Aldrich | CAS 6101-15-1 |
| Paraformaldehyde (PFA) | Sigma-Aldrich | CAS 30525-89-4 |
| Agarose | Sigma-Aldrich | CAS 9012-36-6 |
| Donkey Serum | Merck Millipore | S30-100ML |
| Triton X-100 | Sigma-Aldrich | CAS 9002-93-1 |
| Primary anti-Fos antibody | Synaptic Systems | N/A |
| Secondary anti-rabbit antibody (Alexa Fluor TM 647 coupled) | Invitrogen | N/A |
| DAPI | Sigma-Aldrich | CAS 28718-90-3 |
| Mounting Medium (VectaShield Antifade) | Vector Laboratories | N/A |
| **Deposited data** |  |  |
| All mouse data and code used to generate manuscript figures | This paper | zenodo.org |
| Patient data | This paper | On request |
| **Experimental models: Organisms/strains** |  |  |
| *Mus musculus:* C57BL/6 | Charles River | N/A |
| *Mus musculus:* 129S4-Plxnd1tm1.1(cre/ERT2)Zjh | Jackson Laboratories | RRID:IMSR_JAX:036294 |
| **Software and algorithms** |  |  |
| MATLAB (2023b) | The MathWorks | RRID:SCR_001622 |
| LabVIEW | National Instruments | RRID:SCR_014325 |
| Python | python.org | RRID:SCR_008394 |

**Table S1.** (statistics table) All values are rounded to two significant figures. The tests used were two-tailed hierarchical bootstrap, two-tailed bootstrap, one-tailed bootstrap, F-test, and t-test. Benjamini-Hochberg correction was used for false discovery rate (FDR) adjustment in multiple comparisons, in which case the adjusted p-value p_adj_ is the one reported in the associated figure and text. Sample sizes were estimated based on previous experiments performed in mouse behavioral paradigms (Attinger et al., 2017; Heindorf et al., 2018).

| ***Figure*** | **Description** | **Test** | **N<sub>1</sub>** | **N<sub>2</sub>** | **Unit** | **p-value<br>(p<sub>adj</sub>)** |
|---|---|---|---|---|---|---|
| ***1G*** | Phase III activity duration in mouse EEG (N<sub>1</sub>) vs patient EEG (N<sub>2</sub>) | Two-tailed hierarchical bootstrap | 402 (16) | 50 (26) | Electrodes (mice) vs electrodes (patients) | 0.34 |
| ***1H*** | Power-weighted mean frequency |  |  |  |  |  |
|  | Mouse WF peak (N<sub>1</sub>) vs offset (N<sub>2</sub>) | Two-tailed hierarchical bootstrap, FDR adjusted | 402 (16) | 402 (16) | ROIs (mice) | 0.0014 (0.0032) |
|  | Mouse EEG peak (N<sub>1</sub>) vs offset (N<sub>2</sub>) | Two-tailed hierarchical bootstrap, FDR adjusted | 402 (16) | 402 (16) | Electrodes (mice) | < 0.0010 (< 0.0010) |
|  | Patient EEG peak (N<sub>1</sub>) vs offset (N<sub>2</sub>) | Two-tailed hierarchical bootstrap, FDR adjusted | 50 (26) | 50 (26) | Electrodes (patients) | < 0.0010 (< 0.0010) |
|  | Mouse WF peak (N<sub>1</sub>) vs mouse EEG peak (N<sub>2</sub>) | Two-tailed hierarchical bootstrap, FDR adjusted | 402 (16) | 402 (16) | ROIs (mice) vs electrodes (mice) | 0.025 (0.034) |
|  | Mouse WF offset (N<sub>1</sub>) vs mouse EEG offset (N<sub>2</sub>) | Two-tailed hierarchical bootstrap, FDR adjusted | 402 (16) | 402 (16) | ROIs (mice) vs electrodes (mice) | 0.0018 (0.0032) |
|  | Mouse EEG peak (N<sub>1</sub>) vs patient EEG peak (N<sub>2</sub>) | Two-tailed hierarchical bootstrap, FDR adjusted | 402 (16) | 50 (26) | Electrodes (mice) vs electrodes (patients) | 0.092 (0.11) |
|  | Mouse EEG offset (N<sub>1</sub>) vs patient EEG offset (N<sub>2</sub>) | Two-tailed hierarchical bootstrap, FDR adjusted | 402 (16) | 50 (26) | Electrodes (mice) vs electrodes (patients) | 0.29 (0.29) |
| ***2D*** | Maximum ΔF/F0 in dorsal cortex |  |  |  |  |  |
|  | Pre-ECS (N<sub>1</sub>) vs direct ECS response (N<sub>2</sub>) | Two-tailed hierarchical bootstrap, FDR adjusted | 72 (28) | 72 (28) | ECS (mice) | < 0.0010 (< 0.0010) |
|  | Pre-ECS (N<sub>1</sub>) vs calcium event (N<sub>2</sub>) | Two-tailed hierarchical bootstrap, FDR adjusted | 72 (28) | 22 (18) | ECS (mice) | < 0.0010 (< 0.0010) |
|  | Direct ECS response (N<sub>1</sub>) vs calcium event (N<sub>2</sub>) | Two-tailed hierarchical bootstrap, FDR adjusted | 72 (28) | 22 (18) | ECS (mice) | < 0.0010 (< 0.0010) |
| ***3F*** | Linear regression of ECS charge against probability of calcium event | F-test | 1 | 14 | Degrees of freedom vs Degrees of Freedom for Error | < 0.0010 |
| ***3G*** | Linear regression of ECS charge against probability of phase III activity | F-test | 1 | 14 | Degrees of freedom vs Degrees of Freedom for Error | 0.049 |
| ***4E*** | Mean ΔF/F0 during calcium event in superficial layers somata (N<sub>1</sub>) vs deep layers somata (N<sub>2</sub>) | Two-tailed bootstrap | 236 | 134 | ROIs | see Figure |
| ***4F*** | Time of 50% max. ΔF/F0 in superficial layers somata (N<sub>1</sub>) vs deep layers somata (N<sub>2</sub>) | Two-tailed hierarchical bootstrap | 236 (3) | 134 (3) | ROIs (mice) | < 0.0010 |
| ***4H*** | Mean ΔF/F0 during calcium event in somata (N<sub>1</sub>) vs neuropil (N<sub>2</sub>) | Two-tailed bootstrap | 858 | 858 | ROIs | see Figure |
| ***4I*** | Slope ΔF/F0 of somata (N<sub>1</sub>) vs neuropil (N<sub>2</sub>) | Two-tailed hierarchical bootstrap | 858 (6) | 858 (6) | ROIs (mice) | 0.002 |
| ***5C*** | Fos energy fold-change vs. Sham ECS |  |  |  |  |  |
|  | Sham (N<sub>1</sub>) vs no calcium event (N<sub>2</sub>) | Two-tailed hierarchical bootstrap, FDR adjusted | 10 (5) | 14 (10) | Hemispheres (mice) | 0.58 (0.58) |
|  | No calcium event (N<sub>1</sub>) vs calcium event (N<sub>2</sub>) | Two-tailed hierarchical bootstrap, FDR adjusted | 14 (10) | 10 (8) | Hemispheres (mice) | < 0.0010 (< 0.0010) |
|  | Sham (N<sub>1</sub>) vs calcium event (N<sub>2</sub>) | Two-tailed hierarchical bootstrap, FDR adjusted | 10 (5) | 10 (8) | Hemispheres (mice) | < 0.0010 (< 0.0010) |
| ***6C*** | Mid-phase III activity amplitude |  |  |  |  |  |
|  | Direct ECS response (N<sub>1</sub>) vs contra. calcium event (N<sub>2</sub>) | Two-tailed hierarchical bootstrap, FDR adjusted | 93 (4) | 52 (4) | Electrodes (mice) | 0.34 (0.34) |
|  | Direct ECS response (N<sub>1</sub>) vs ipsi. calcium event (N<sub>2</sub>) | Two-tailed hierarchical bootstrap, FDR adjusted | 93 (4) | 257 (12) | Electrodes (mice) | < 0.0010 (< 0.0010) |
|  | Contra. calcium event (N<sub>1</sub>) vs ipsi. calcium event (N<sub>2</sub>) | Two-tailed hierarchical bootstrap, FDR adjusted | 52 (4) | 257 (12) | Electrodes (mice) | 0.016 (0.025) |
| ***6D*** | Phase III activity duration |  |  |  |  |  |
|  | Direct ECS response (N<sub>1</sub>) vs contra. calcium event (N<sub>2</sub>) | Two-tailed hierarchical bootstrap, FDR adjusted | 93 (4) | 52 (4) | Electrodes (mice) | 0.33 (0.50) |
|  | Direct ECS response (N<sub>1</sub>) vs ipsi. calcium event (N<sub>2</sub>) | Two-tailed hierarchical bootstrap, FDR adjusted | 93 (4) | 257 (12) | Electrodes (mice) | 0.00040 (0.0012) |
|  | Contra. calcium event (N<sub>1</sub>) vs ipsi. calcium event (N<sub>2</sub>) | Two-tailed hierarchical bootstrap, FDR adjusted | 52 (4) | 257 (12) | Electrodes (mice) | 0.52 (0.52) |
| ***6E*** | Postictal suppression index |  |  |  |  |  |
|  | Direct ECS response (N<sub>1</sub>) vs contra. calcium event (N<sub>2</sub>) | Two-tailed hierarchical bootstrap, FDR adjusted | 93 (4) | 52 (4) | Electrodes (mice) | 0.13 (0.19) |
|  | Direct ECS response (N<sub>1</sub>) vs ipsi. calcium event (N<sub>2</sub>) | Two-tailed hierarchical bootstrap, FDR adjusted | 93 (4) | 257 (12) | Electrodes (mice) | < 0.0010 (< 0.0010) |
|  | Contra. calcium event (N<sub>1</sub>) vs ipsi. calcium event (N<sub>2</sub>) | Two-tailed hierarchical bootstrap, FDR adjusted | 52 (4) | 257 (12) | Electrodes (mice) | 0.86 (0.86) |
| ***6F*** | Inter-hemispheric coherence |  |  |  |  |  |
|  | Direct ECS response (N<sub>1</sub>) vs contra. calcium event (N<sub>2</sub>) | Two-tailed hierarchical bootstrap, FDR adjusted | 93 (4) | 52 (4) | Electrodes (mice) | 0.27 (0.61) |
|  | Direct ECS response (N<sub>1</sub>) vs ipsi. calcium event (N<sub>2</sub>) | Two-tailed hierarchical bootstrap, FDR adjusted | 93 (4) | 257 (12) | Electrodes (mice) | 0.41 (0.61) |
|  | Contra. calcium event (N<sub>1</sub>) vs ipsi. calcium event (N<sub>2</sub>) | Two-tailed hierarchical bootstrap, FDR adjusted | 52 (4) | 257 (12) | Electrodes (mice) | 0.70 (0.70) |
| ***S1B*** | Mean ΔF/F0 during ECS vs baseline | Two-tailed bootstrap | 58744 | - | ROIs | see Figure |
| ***S3A*** | Linear regression of ECS parameter against probability of calcium event |  |  |  |  |  |
|  | ECS frequency against probability of calcium event | F-test | 1 | 2 | Degrees of freedom vs Degrees of Freedom for Error | 0.13 |
|  | ECS current against probability of calcium event | F-test | 1 | 2 | Degrees of freedom vs Degrees of Freedom for Error | 0.052 |
|  | Multivariate linear regression of ECS parameter against probability of calcium event |  |  |  |  |  |
|  | ECS frequency against probability of calcium event | Two-sided t-test | 1 | 12 | Degrees of freedom vs Degrees of Freedom for Error | 0.76 |
|  | ECS current against probability of calcium event | Two-sided t-test | 1 | 12 | Degrees of freedom vs Degrees of Freedom for Error | 0.43 |
|  | Interaction of ECS frequency and ECS current against probability of calcium event | Two-sided t-test | 1 | 12 | Degrees of freedom vs Degrees of Freedom for Error | 0.019 |
| ***S3B*** | Linear regression of ECS parameter against probability of phase III activity |  |  |  |  |  |
|  | ECS frequency against probability of phase III activity | F-test | 1 | 2 | Degrees of freedom vs Degrees of Freedom for Error | 0.14 |
|  | ECS current against probability of phase III activity | F-test | 1 | 2 | Degrees of freedom vs Degrees of Freedom for Error | 0.37 |
| ***S4A*** | Linear regression of ECS charge against maximum ΔF/F0 during direct ECS response | F-test | 1 | 70 | Degrees of freedom vs Degrees of Freedom for Error | < 0.0010 |
| ***S4B*** | Maximum ΔF/F0 during direct ECS response in ipsilateral (N<sub>1</sub>) over contralateral hemisphere (N<sub>2</sub>) | Two-tailed hierarchical bootstrap | 16 (13) | 16 (13) | ECS (mice) | 0.0086 |
| ***S5A*** | Mean ΔF/F0 during ECS |  |  |  |  |  |
|  | EGFP (N1) vs baseline | Two-tailed bootstrap | 14190 | - | ROIs | see Figure |
|  | EGFP (N1) vs GCaMP6s (N<sub>2</sub>) | Two-tailed bootstrap | 14190 | 58744 | ROIs | see Figure |
| ***S5C*** | Maximum ΔF/F0 in dorsal cortex during calcium event (N<sub>1</sub>) vs EGFP event (N<sub>2</sub>) | Two-tailed hierarchical bootstrap | 22 (18) | 8 (6) | ECS (mice) | < 0.0010 |
| ***S5D*** | Propagation speed of calcium event (N<sub>1</sub>) vs EGFP event (N<sub>2</sub>) | Two-tailed hierarchical bootstrap | 22 (18) | 8 (6) | ECS (mice) | 0.10 |

**Table S2.**
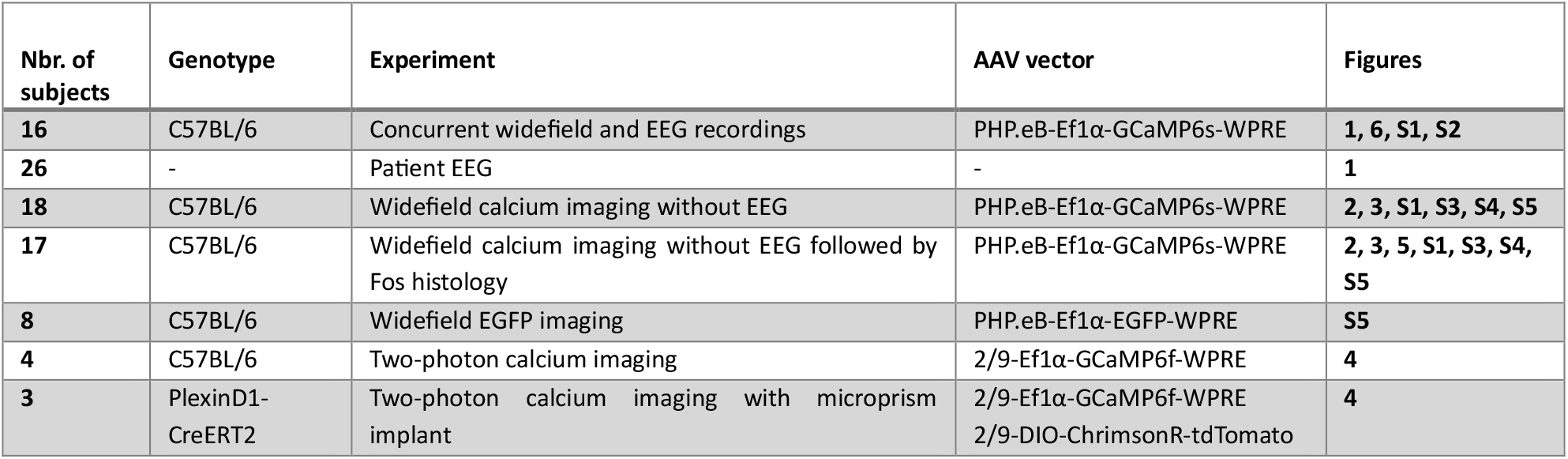
Number of subjects in each analysis.

| **Nbr. of subjects** | **Genotype** | **Experiment** | **AAV vector** | **Figures** |
|---|---|---|---|---|
| **16** | C57BL/6 | Concurrent widefield and EEG recordings | PHP.eB-Ef1α-GCaMP6s-WPRE | **1, 6, S1, S2** |
| **26** | - | Patient EEG | - | **1** |
| **18** | C57BL/6 | Widefield calcium imaging without EEG | PHP.eB-Ef1α-GCaMP6s-WPRE | **2, 3, S1, S3, S4, S5** |
| **17** | C57BL/6 | Widefield calcium imaging without EEG followed by Fos histology | PHP.eB-Ef1α-GCaMP6s-WPRE | **2, 3, 5, S1, S3, S4, S5** |
| **8** | C57BL/6 | Widefield EGFP imaging | PHP.eB-Ef1α-EGFP-WPRE | **S5** |
| **4** | C57BL/6 | Two-photon calcium imaging | 2/9-Ef1α-GCaMP6f-WPRE | **4** |
| **3** | PlexinD1-CreERT2 | Two-photon calcium imaging with microprism implant | 2/9-Ef1α-GCaMP6f-WPRE<br>2/9-DIO-ChrimsonR-tdTomato | **4** |

**Table S3.** Distribution of stimulation parameters used for ECS in mice. Stimulation duration and pulse width were kept constant at 1 s and 0.5 ms, respectively (**Methods**).

| **Nbr. of ECS** | **Stimulation frequency [Hz]** | **Stimulation amplitude [mA]** | **Single ECS example** | **Figures** |
|---|---|---|---|---|
| **26** | 100 | 55 | **2F, 5A (left and middle row), 5B (middle and right row), S5A, S5B, Video S1** | **1G, 1H, 2C, 2D, 3A, 3B, 3C, 3D, 3E, 3F, 3G, 5C, 6C, 6D, 6E, 6F, S1B, S3A, S3B, S3C, S3D, S3E, S4A, S4B, S5C, S5D** |
| **28** | 100 | 75 | **1B, 1C, 1F, 4C, 4D, 5B (left row), 6A, 6B, S1C, S2, Video S2** | **1G, 1H, 2C, 2D, 3A, 3B, 3C, 3D, 3E, 3F, 3G, 4E, 4F, 4H, 4I, 4J, 5C, 6C, 6D, 6E, 6F, S1B, S3A, S3B, S3C, S3D, S3E, S4A, S4B, S5C, S5D** |
| **19** | 100 | 35 | **Video S1** | **1G, 1H, 2C, 2D, 3A, 3B, 3C, 3D, 3E, 3F, 3G, 4H, 4I, 4J, 6C, 6D, 6E, 6F, S1B, S3A, S3B, S3C, S3D, S3E, S4A, S4B, S5C, S5D** |
| **9** | 50 | 35 |  | **2D, 3C, 3D, 3E, 3F, 3G, S1B, S3A, S3B, S4A** |
| **8** | 25 | 75 |  | **1G, 1H, 2C, 2D, 3A, 3B, 3C, 3D, 3E, 3F, 3G, 6C, 6D, 6E, 6F, S1B, S3A, S3B, S3C, S3D, S3E, S4A, S4B** |
| **8** | 50 | 55 | **S1A** | **2D, 3C, 3D, 3E, 3F, 3G, S1B, S3A, S3B, S4A** |
| **7** | 75 | 35 |  | **2C, 2D, 3A, 3B, 3C, 3D, 3E, 3F, 3G, S1B, S3A, S3B, S3C, S3D, S3E, S4A, S4B** |
| **6** | 75 | 75 | **2G, 4B, Video S3** | **1G, 1H, 2C, 2D, 3A, 3B, 3C, 3D, 3E, 3F, 3G, 4H, 4I, 4J, 6C, 6D, 6E, 6F, S1B, S3A, S3B, S3C, S3D, S3E, S4A, S4B** |
| **5** | 25 | 15 |  | **2D, 3C, 3D, 3E, 3F, 3G, S1B, S3A, S3B, S4A** |
| **5** | 75 | 15 |  | **2D, 3C, 3D, 3E, 3F, 3G, 4H, 4I, 4J, S1B, S3A, S3B, S4A** |
| **5** | 75 | 55 |  | **2D, 3C, 3D, 3E, 3F, 3G, S1B, S3A, S3B, S4A** |
| **4** | 50 | 15 |  | **2D, 3C, 3D, 3E, 3F, 3G, S1B, S3A, S3B, S4A** |
| **4** | 100 | 15 |  | **2D, 3C, 3D, 3E, 3F, 3G, S4A, S3A, S3B, S1B, 4H, 4I, 4J** |
| **3** | 25 | 35 |  | **2D, 3C, 3D, 3E, 3F, 3G, S1B, S3A, S3B, S4A** |
| **3** | 25 | 55 |  | **2D, 3C, 3D, 3E, 3F, 3G, S1B, S3A, S3B, S4A** |
| **3** | 50 | 75 |  | **2D, 3C, 3D, 3E, 3F, 3G, S1B, S3A, S3B, S4A** |
| **3** | 100 | 20 |  | **2D, 3C, 3D, 3E, 3F, 3G, 5C, S1B, S3A, S3B, S4A** |
| **3** | 100 | 50 | **2B, 2E, 5A (right row), Video S1** | **2C, 2D, 3A, 3B, 3C, 3D, 3E, 3F, 3G, 5C, S3A, S3B, S3C, S3D, S3E, S4A, S4B** |

**Table S4.** Stimulation parameters used for ECT in patients. The example patient ECT data shown in **Figure 1E-F** is from patient no. 12.

| Patient no. | Stimulation frequency [Hz] | Stimulation amplitude [mA] | Stimulation duration [s] | Pulse width [ms] |
|---|---|---|---|---|
| 1 | 70 | 910 | 6.9 | 0.75 |
| 2 | 40 | 910 | 6.3 | 0.5 |
| 3 | 60 | 910 | 7.5 | 0.75 |
| 4 | 60 | 900 | 7.5 | 0.75 |
| 5 | 40 | 910 | 7.7 | 0.5 |
| 6 | 70 | 910 | 7.5 | 0.75 |
| 7 | 70 | 910 | 6.9 | 0.75 |
| 8 | 40 | 910 | 7 | 0.5 |
| 9 | 60 | 900 | 7.9 | 0.5 |
| 10 | 60 | 900 | 7.5 | 0.75 |
| 11 | 70 | 900 | 7.6 | 0.5 |
| 12 | 70 | 900 | 8 | 0.5 |
| 13 | 50 | 910 | 6.7 | 0.5 |
| 14 | 40 | 920 | 7 | 0.5 |
| 15 | 40 | 910 | 7.7 | 0.5 |
| 16 | 40 | 900 | 6.3 | 0.5 |
| 17 | 60 | 890 | 7.5 | 0.5 |
| 18 | 50 | 900 | 7.8 | 0.5 |
| 19 | 50 | 900 | 6.7 | 0.5 |
| 20 | 50 | 910 | 7.8 | 0.5 |
| 21 | 60 | 900 | 6.8 | 0.75 |
| 22 | 50 | 900 | 7.8 | 0.5 |
| 23 | 70 | 910 | 7.5 | 0.75 |
| 24 | 50 | 910 | 7.8 | 0.5 |
| 25 | 70 | 900 | 8 | 0.5 |
| 26 | 50 | 910 | 6.7 | 0.5 |

